# Spillover rate is not a predictor of host jump risk, but spillover novelty is

**DOI:** 10.1101/2024.10.14.618291

**Authors:** Brandon J. Simony, David A. Kennedy

**Author notes:** B.J.S. and D.A.K. conceptualized the project, developed the model and methodology, and revised and edited the manuscript. B.J.S. wrote the original draft of the manuscript, wrote the software, conducted the formal analysis, validated the results, and generated the figures. D.A.K. acquired the funding, provided mentorship, and oversaw the project. The authors declare no competing interests.

## Abstract

Host jumps—defined as the process by which a pathogen establishes sustained transmission in novel hosts—are threats to human and animal welfare, but prediction of successful host jumps remains elusive. A spillover event must precede a host jump, and so spillover rate is thought to be related to risk. However, non-endemic pathogens that spill over frequently have demonstrated a poor ability to host jump from any given spillover. So which is riskier, pathogens that spill over rarely or commonly? Applying a Bayesian framework to a general model of host jump risk, we show that 1) the riskiest pathogens can be those that spill over at low, intermediate, or high rates, and 2) as the rate of spillover gets large, the information gained from past spillovers is exactly counterbalanced by the increased number of future spillovers. Taken together, this means that spillover rate has little to no value for explaining host jump risk. Rather, we show that novel pathogens (i.e. pathogens with a relatively short history of spilling over in their current form) are substantially more likely to result in host jumps than pathogens that have had long-associated opportunities for spillover into the novel host. Therefore novelty, but not spillover rate, is an important predictor of host jump risk.

**Significance Statement:** Spillovers of infectious diseases pose significant threats to human and animal health in part due to pandemic risk and the potential for successful host jumps. Predicting successful host jumps remains elusive, though it is often assumed that increased spillover rates are associated with increased host jump risk. We developed a quantitative framework that uses past spillover outcomes to predict future host jump risk. Our model demonstrates that spillover rate is not predictive of host jump risk. However, pathogens that recently started having opportunities for spillover are substantially riskier than those that have a longer history of past spillover. We therefore show that pathogen novelty is a much stronger driver of host jump risk than spillover rate. This ideological shift may significantly enhance our ability to identify contexts where host jumps are most likely, yielding substantial economic and public health benefits globally.

---

Zoonotic pathogens—defined as animal pathogens that can spill over into humans—are a growing concern to global public health, and it is expected that they will be responsible for the next pandemic (1, 2). Many recognized human pathogens are believed to be of zoonotic origin (3–5), and a large body of research has focused on identifying characteristics that predispose a pathogen to successfully establish itself and persist in a new host species, hereafter referred to as a host jump (5–15). Spillover—defined as the events during which a pathogen transmits from a native host species into a novel host species—is a prerequisite to a host jump, and so efforts to reduce host jump risk have largely focused on pathogens that spill over frequently (16–20). However, it is unclear how a pathogen’s inherent rate of spillover relates to the likelihood of a successful host jump. Put another way, are pathogens that frequently spill over more or less likely to host jump than those that rarely spill over?

At present, the answer to the aforementioned question is unknown, and the risk of a host jump undoubtedly depends on many system-specific details, such as characteristics of the native and novel hosts, the set of all zoonotic pathogens, and other ecological and evolutionary hurdles. However, by considering two pathogens that differ only in their inherent rate of spillover, we can evaluate how spillover rate itself impacts host jump risk. On one hand, a pathogen that frequently spills over has many opportunities to complete a host jump, so all else equal, a pathogen that spills over more frequently may be conceivably more likely to complete a host jump. On the other hand, a pathogen that has not yet jumped hosts despite having spilled over many times has demonstrated an inability to overcome the ecological and evolutionary hurdles necessary to successfully host jump (18, 21), and so pathogens that have spilled over at high rates in the past may be less likely to host jump. Whether a pathogen with a high spillover rate is more likely to host jump than a pathogen with a low spillover rate depends on how these two competing factors combine together.

To further illustrate the interplay between spillover rate and host jump likelihood, consider several examples. Rabies lyssavirus, the causative agent of rabies, is estimated to cause tens of thousands of human rabies cases per year due to zoonotic spillover from dogs and other animal reservoirs (22, 23). Rabies presents a significant public health burden, so preventive measures in place to mitigate spillover are undoubtedly important (24). However, even without these preventative measures and despite the high rate of spillover, few would argue that rabies is a pandemic threat, given the limited opportunities for human-to-human transmission, which would presumably require a human biting another human (25). A similar argument could be made for *Borellia burgdorferi*, the causative agent of Lyme disease, where sustained human transmission would be limited by the frequency and duration of ticks feeding on humans (26). In contrast, human immunodeficiency virus (HIV) has completed host jumps into humans on at least two separate occasions (27), and these host jumps occurred after a presumably smaller number of spillover events than the aforementioned pathogens. Despite being a hand-picked list, these examples do illustrate the potential for a disconnect between spillover rate and host jump risk.

Here, we develop a model that provides a quantitative framework for evaluating host jump risk. We derive an analytical solution to this model that depends on the rate of past and future spillover events, the duration of opportunities for spillover in the past, the time window over which host jump risk is being assessed, and our uncertainty regarding the probability that a single spillover event will result in a host jump. Using this analytical solution, we show that a pathogen’s host jump risk is highly insensitive to the pathogen’s inherent rate of spillover. Rather, the key predictor of host jump risk is “pathogen novelty” (defined as pathogens that either did not have opportunities for spillover in the past, or that have evolved suffciently far that the outcome of past spillover events is unrelated to the outcome of future spillover events.

## 1. Model Overview

It is worth acknowledging that despite all the complexities of biology, a successful host jump can be exactly distilled into a simple two-part process. First, a pathogen must spill over into a novel host thereby infecting it, and second, the pathogen must spread suffciently well as to establish sustained transmission in the novel host population. To complete a host jump, pathogens must overcome barriers in both of these processes (20), and pathogens can therefore be roughly divided into four categories: “unrestricted”, “transmission limited”, “spillover limited”, and “spillover and transmission limited” (Figure 1A-D). We define “unrestricted pathogens” as those that frequently spill over and are also likely to sustain transmission in the novel host population. “Transmission limited” pathogens are those that frequently spill over but are unlikely to sustain transmission in the novel host. “Spillover limited” pathogens are those that rarely spill over but are likely to sustain transmission in the novel host if they do. “Spillover and transmission limited” pathogens are those that rarely spill over and are also unlikely to sustain transmission in the novel host population.

**Fig. 1.**
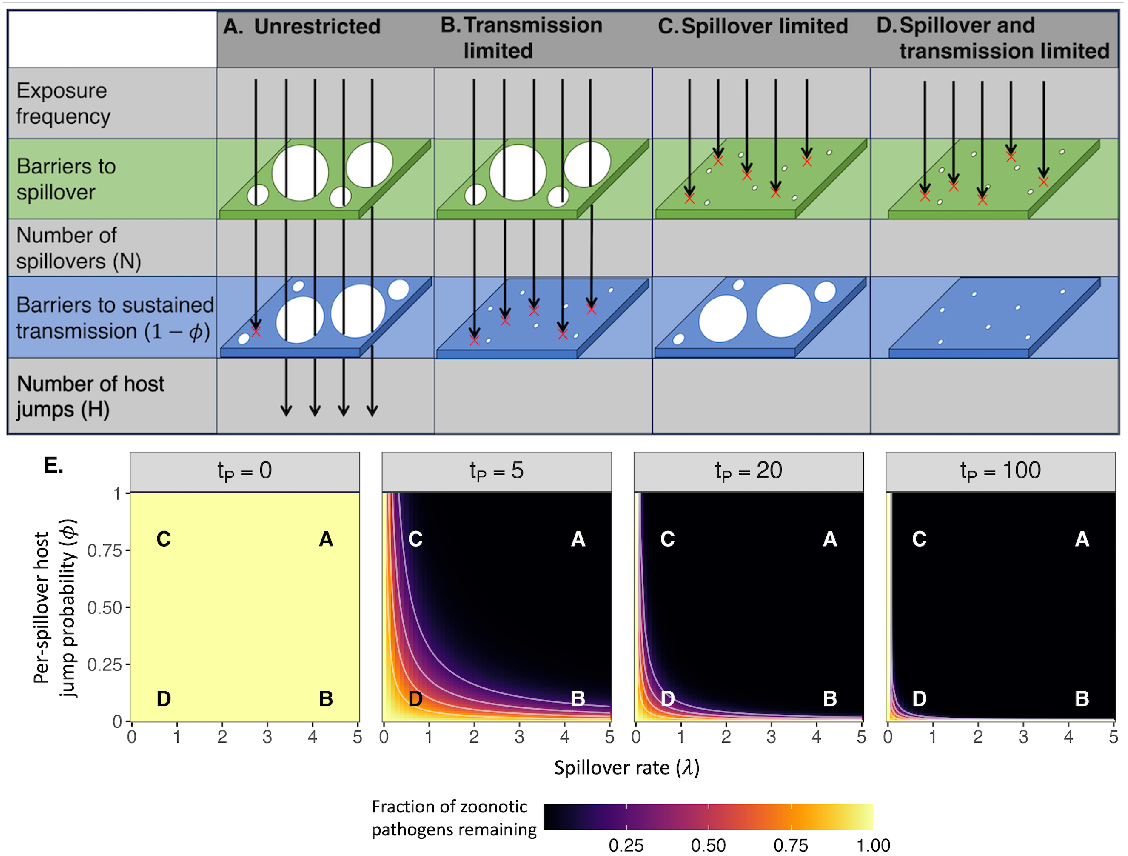
Conceptual characterization of possible pathogen types (**A-D**). In order to successfully host jump, a pathogen must overcome barriers to spillover to successfully infect a novel host (20). Following a spillover event, the pathogen must also be able to overcome the ecological and evolutionary barriers to sustained transmission in the novel host (21). Pathogens may or may not be limited at either step in this process, and the composition of the pool of zoonotic pathogens likely changes over time. In panel **E**, we show how the pool of zoonotic pathogens may change as a function of the duration of past opportunities for spillover *t*_*P*_. Note that *t*_*P*_ is not a measure of absolute time, but rather relative time, and will therefore differ among zoonotic pathogens depending on how long each pathogen had opportunities for spillover. For illustrative purposes, we assume that at the initial time point (*t*_*P*_ = 0), all pathogens are equally frequent. This panel demonstrates that as *t*_*P*_ increases, the proportion of “unrestricted” zoonotic pathogens (**A**) decreases quickly, meaning that as *t*_*P*_ gets larger, pathogens are increasingly likely to be transmission limited (**B**) spillover limited (**C**) or both (**D**).

The proportion of pathogens that fall within each of these categories is likely to change over long periods of time, in part because once a host jump occurs, the pathogen that jumped hosts changes from being a zoonotic pathogen to being a human pathogen of zoonotic origin. Figure 1E illustrates how this composition changes over time *t*_*P*_as a function of how spillover-limited and transmission-limited pathogens are. Here *t*_*P*_is the difference between the current time and the time when a pathogen first arose and had opportunities to spill over. Note that we can quickly become confident that few unrestricted zoonotic pathogens remain in the zoonotic pool because any that exist would quickly complete host jumps. Thus at any given time, most zoonotic pathogens will be spillover limited, transmission limited, or both. Note, however, that *t*_*P*_will differ for different pathogens depending on how long each had opportunities for spillover in its current form.

To develop a quantitative framework for host jump risk, it is first useful to realize that there are only two possible outcomes that can occur following a spillover event; either the pathogen dies out in the novel host population or it establishes itself. Although the precise disease dynamics that arise following a spillover event will depend on a complicated demographic process, the final outcome of a given spillover event (i.e. host jump or not) is simply a Bernoulli trial with a probability that is, in theory, determinable from that complicated demographic process. Using this fact, we can quantify the probability that no host jump has occurred in some number of spillover events (*N*). To do so, we define *ϕ* as the fixed probability that any given spillover event results in a successful host jump, and we assume the outcome of individual spillovers are independent. Since the only possible outcomes of a single spillover event are a host jump or not, we can quantify the probability that no host jump occurs (*H* = 0) with a binomial distribution such that *P* (*H* = 0*|ϕ, N*) = (1 −*ϕ*)^*N*^. As an aside, each spillover event presumably has a unique host jump probability (*ϕ*_*j*_) that depends on interactions between features of the host, pathogen, and environment in which the spillover event occurred. However, our equation is unchanged even in the presence of this variation, provided *ϕ* is set to the average of *ϕ*_*j*_, and the average of *ϕ*_*j*_does not change over time (Supplementary text A).

Moving from counts of spillover events to rates of spillover events, we can analogously define the probability that no host jump occurs within a finite time *t* using a Poisson process with parameter *λt*, where *λ* is the pathogen’s rate of spillover (see Supplementary text B) and *t* is the time interval over which the probability of a host jump is being considered. This yields the probability that no host jump occurs within a time interval of duration *t*,

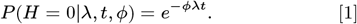

Thus, when the outcome of each spillover event is independent, Eq. 1 exactly describes the probability that a zoonotic pathogen will not successfully host jump within a particular time window. Note that this math is true irrespective of the biological complexities of the system, with the exception of the assumptions already laid out. One of the key challenges with using this equation, however is that *ϕ* (the probability of persistence for a single spillover event) is typically unknown. Here, we propose that the past can help inform the value of *ϕ*. To illustrate our logic, consider a pathogen that has spilled over millions of times in the past, but has not yet completed a host jump. Intuitively, its value of *ϕ* must be very small, since it likely would have already jumped hosts otherwise.

This logic can be formalized by employing tools from Bayesian statistics, in which past observations of spillover that failed to result in a host jump update our belief in the value of a pathogen’s ability to successfully host jump following a spillover event (i.e. *ϕ*). To do so, we treat *ϕ* as a distribution rather than a fixed constant, where *π*(*ϕ*) is the distribution that describes our initial uncertainty regarding the value of *ϕ* (in Bayesian statistics, this is referred to as a prior). One advantage of this approach is that it explicitly allows our uncertainty to be updated in a statistically rigorous way as new data become available. Following the standard definition of Bayes’ theorem, our posterior distribution of *ϕ* is simply

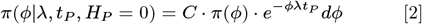

where *C* is a normalizing constant and the subscript *P* denotes that parameters are referring to the past.

We can then use this posterior distribution of *ϕ* to quantify future host jump risk. The probability of no host jumps occurring in some future time interval of duration *t*_*F*_can be expressed as the integrated product of Eq. 2 and Eq. 1. Therefore, one minus this quantity gives the probability that, for a given set of pathogens, at least one future spillover event will result in a successful host jump, as shown in equation 3. More precisely:

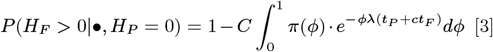

This value is conditional on the model parameters (represented by *•*) for the rates of past and future spillover (*λ* and *cλ* respectively), the duration of opportunities for past spillover (*t*_*P*_), the future time horizon (*t*_*F*_), and our prior belief about the likelihood that a given spillover event will result in a host jump *π*(*ϕ*). In the present analyses, we only consider cases where no past spillover event resulted in a host jump (i.e., *H*_*P*_= 0), but this equation can be readily extended—although perhaps less usefully—to cases where past spillovers have resulted in successful host jumps (Supplementary text B).

To finish parameterizing our model, it is necessary to define *π*(*ϕ*), which reflects our initial uncertainty regarding the probability that a single spillover event will result in a host jump. However, specifying an appropriate shape for this distribution is not straightforward and may differ for different groups of pathogens. We therefore consider three shapes for this distribution:

1. The vast majority of zoonotic pathogens are unlikely to persist in the novel host, meaning the distribution of *ϕ* is extremely right-skewed (Fig. 2A).
2. Zoonotic pathogens fall into two distinct sub-classes, where some pathogens are highly likely to persist in the novel host, and the rest are unlikely to persist, meaning that the distribution of *ϕ* is “U-shaped” (Fig. 2B).
3. Most zoonotic pathogens are unlikely to persist in the novel host, but a subset have an intermediate chance of persisting, meaning the distribution of *ϕ* is multi-modal with one peak at 0 and another peak at an intermediate value (Fig. 2C).

**Fig. 2.**
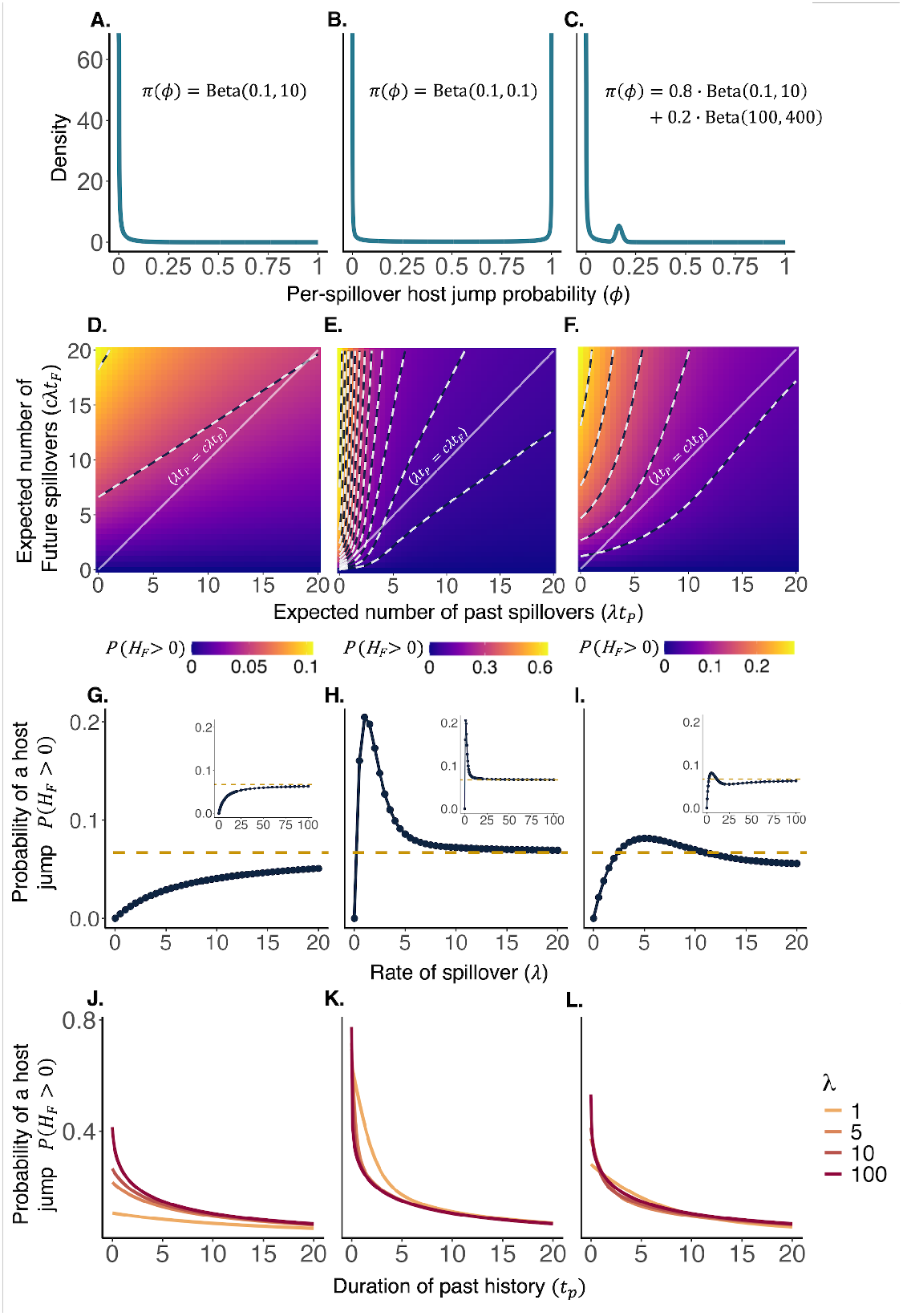
The columns of panels from left to right denote Scenarios 1-3 respectively. Panels A-C show the prior distributions on *ϕ*. Panels D-F show the model-calculated probability of a successful host jump as determined from Eq. 4. Axes show the effects of changing the two key parameter combinations, with the x-axes showing the effect of varying the composite parameter of past spillover rate *λ* times the duration of opportunities for spillover *t*_*P*_, and the y-axes showing the effects of varying the composite parameter of future spillover rate *cλ* times the time window over which host jump risk is being assessed *t*_*F*_. The contour lines in each panel demarcate changes of 5% in total host jump risk. Note that the color range differs for each panel. The solid white lines depict a 1:1 line, which can arise when the rate of spillover in the past and future are equal. Panels G-I show the probabilities of a future host jump along these 1:1 lines. Comparing panels G-I reveals that depending on the prior *π*(*ϕ*), host jump risk may be greatest at high rates of spillover, low rates of spillover or intermediate rates of spillover. Nevertheless, in all scenarios, host jump risk converges to the value 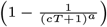 (Eq. 5), as spillover rate increases (inset figures) demonstrating that as a pathogen’s inherent rate of spillover increases, the rate gives progressively less information about host jump risk. In panels J-L, we plot the probability of a future host jump as a function of the duration of time over which spillover opportunities existed in the past (*t*_*P*_). In each scenario, the probability of a host jump declines monotonically and dramatically as *t*_*P*_ increases from zero, demonstrating that the greatest risk of a host jump is posed by pathogens that are new. Furthermore, these panels show that even with drastically different rates of spillover, host jump risk is much more strongly impacted by changes in the duration of past history *t*_*P*_ than changes in the rate of spillover *λ*, adding support to our claim that spillover rate is not a useful predictor of host jump risk.

Scenario 1 and 2 above can be modeled using a beta distribution and Scenario 3 can be modeled using a mixture of two beta distributions. The results of our analyses are general, but for illustration purposes, we will specify parameter values for each of our prior distributions such that in Scenario 1: *π*(*ϕ*) ~ Beta(*a* = 0.1, *b* = 10), in Scenario 2: *π*(*ϕ*) ~ Beta(*a* = 0.1, *b* = 0.1), and in Scenario 3: *π*(*ϕ*) ~ *ω*_1_Beta(*a* = 0.1, *b* = 10) + *ω*_2_Beta(*a* = 100, *b* = 400); *ω*_1_ = 0.8, *ω*_2_ = 0.2. In Supplementary text B, we show that when *π*(*ϕ*) follows a beta distribution, there is an analytical solution to Eq. 3 such that:

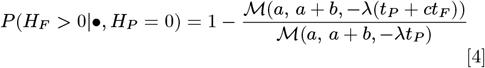

where *•* again represents the set of model parameters and *ℳ* (*a, b, z*) is the confluent hypergeometric function of the first kind (28). We show that a similar analytical solution exists for beta-mixture distributions (Supplementary text D). We also derive an analogous version of this model that uses numbers of spillover events instead of rates showing that our results are fundamentally similar (Supplementary text C).

Note that in Eq. 4 the risk of a host jump depends on a total of six parameters, but only on four parameter combinations: the hyperparameters *a* and *b* of *π*(*ϕ*) (which we explore both in the Supplemental text G and using scenarios 1-3), the past spillover rate *λ* times the duration of time during which spillover was possible in the past *t*_*P*_, and the future spillover rate *cλ* times the future time window over which host jump risk is being assessed *t*_*F*_. In what follows, we use Eq. 4 and its beta-mixture analog to show that host jump risk is largely insensitive to spillover rate *λ* but is highly sensitive to the length of time that spillover was possible in the past *t*_*P*_, thus demonstrating that novelty, rather than spillover rate, is the key driver of host jump risk. All main results were also confirmed using simulation (Supplementary text F).

## Results

Using equation 4, we evaluate the probability of a successful host jump in scenarios 1-3 (Fig. 2 panels D-F respectively) as a function of the key parameter combinations *λt_P_*(expected number of past spillovers) and *cλt_F_*(expected number of future spillovers). In all three scenarios, when only varying the future values, the probability of a future host jump increases as the expected number of future spillovers increases *cλt_F_*(i.e. host jump probability increases when moving from bottom to top in Fig. 2D-F). Intuitively, this should be true, as more spillover events create more opportunities for a host jump to occur. In contrast, when only varying the past values, the probability of a future host jump decreases as the expected number of past spillovers increases (i.e. host jump probability decreases when moving from left to right in Fig. 2D-F). This can be explained by the increased observation of failed host jumps driving the posterior *π*(*ϕ|λ, t_P_, H_P_*= 0) closer to zero, making each future spillover event less likely to result in a host jump. In what follows, we delved deeper into these results to separate the impact of spillover rate *λ* from the impact of time, specifically *t*_*P*_.

With regard to spillover rates, the past and future are often correlated. Pathogens with high past spillover rates are likely to have high future spillover rates, and pathogens with low past spillover rates are likely to have low future spillover rates. We have implicitly assumed this linear relationship by defining the future spillover rate as *cλ*, where *c* is an arbitrary constant that accounts for differences between the past and future due to changes outside the pathogen’s control (we relax the assumption of linearity in Supplemental text D). Using this reformulation, we can then assess the relationship between a pathogen’s inherent spillover rate its risk of a future host jump. Holding *t*_*P*_and *t*_*F*_constant, this means that as a pathogen’s inherent rate of spillover changes, it moves along Fig. 2 panels D-F on a straight line with slope *c*. By comparing host jump risk along this line, we can therefore assess the role of a pathogen’s inherent rate of spillover on its host jump risk.

For illustrative purposes, we show the case where the inherent rate of spillover is equal between the past and future (*c* = 1), but note that all of our results are qualitatively similar for other linear correlations. Figures 2G-I show how the probability of a host jump varies for zoonotic pathogens that are in all ways equivalent except for differences in their inherent rates of spillover. We find that the level of spillover where host jump risk is greatest depends on the prior *π*(*ϕ*). In Scenario 1 (Fig. 2G), increasing spillover leads to a monotonic increase in host jump probability. In Scenario 2 (Fig. 2H), host jump risk peaks at low rates of spillover and thereafter monotonically decreases. In Scenario 3 (Fig. 2I), host jump risk peaks at an intermediate rate of spillover, declines at slightly higher rates of spillover, and then increases again when rates of spillover become very high. These results therefore illustrate that, when comparing across zoonotic pathogens, even with all else equal, pathogens with high rates of spillover may be either more or less likely to cause a host jump than pathogens with low rates of spillover, and the conclusion depends on both our prior uncertainty regarding the value of *ϕ* and the inherent rate of spillover of the pathogens.

Next, we ask how the probability of a host jump changes as the rate of spillover becomes large. To do so, we evaluate the limit of Eq. 4 as *λ* approaches infinity. In Supplementary text D, we show that the probability of a host jump converges to

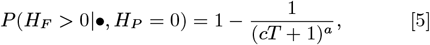

where *a* is the first shape parameter of the beta distribution for prior *π*(*ϕ*), *T* is the ratio between the past and future time intervals 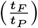, and *c* is the ratio between the past and future spillover rate. Notably, Eq. 5 is strictly between zero and one when *a, c, t_P_*and *t*_*F*_are positive. In Supplemental text D2, we derive the exact same solution for priors that use mixtures of beta distributions, where the value of *a* above is equal to the component of the beta mixture with the smallest *a*. Because the beta shape parameter *a* is the same for the three scenarios used here, all converge to the same value (the dashed gold lines in Fig. 2G-I). However, they can approach this value from below (i.e., higher rates of spillover lead to an increase in host jump risk) or above (i.e., higher rates of spillover lead to a decrease in host jump risk). This implies that our uncertainty regarding the shape of *π*(*ϕ*) can fundamentally change our conclusion as to whether the greatest host jump risk comes from pathogens that spill over frequently or those that spill over rarely. In addition, Fig. 2G-I demonstrates that, except at very low rates of spillover, host jump risk changes very slowly relative to the change in spillover rate, meaning that large differences in spillover rate between two zoonotic pathogens will have little predictive power regarding the overall probability that a host jump will occur.

We next address how the duration of past opportunities for spillover *t*_*P*_relates to the probability of a successful host jump. Note that in contrast to spillover rate, there is no reason to expect *t*_*P*_ to correlate with *t*_*F*_(the time window over which the risk of a future host jump is being assessed). Therefore, in analyzing these results, we assume *t*_*F*_is invariant with regard to *t*_*P*_. Functionally, this is equivalent to moving left or right on Fig. 2D-F. In figure 2J-L, we plot the probability of a future host jump as a function of the duration of past opportunities for spillover *t*_*P*_for each of our three priors. This figure shows that host jump risk is strongly affected by *t*_*P*_, declining rapidly as *t*_*P*_increases. This claim is reinforced by Eq. 5, which shows that the probability of a host jump goes to zero as *t*_*P*_gets large. The figure also reiterates our earlier claim that the rate of spillover only has a minimal effect on host jump risk, and it demonstrates that any variation in host jump risk attributable to differences in spillover rates becomes less important as *t*_*P*_increases. In other words, any two pathogens that had the opportunity to spill over for a similar duration of time *t*_*P*_are roughly equally likely to jump hosts, even if their inherent rates of spillover are dramatically different. Thus, the pathogens at greatest risk of leading to host jumps are not those that spill over frequently, but rather those that are novel, where novelty is defined as zoonotic threats that recently gained the opportunity to spill over. The one caveat here is that for pathogens where *t*_*P*_is exactly equal to 0 (i.e. pathogens that have never yet had an opportunity where spillover was possible), higher rates of spillover are universally worse.

Taken in total, these results suggest that a pathogen’s inherent rate of spillover is not predictive of host jump risk, but rather, spillover novelty is. These results arise for several reasons. First, at low levels of spillover, the choice of *π*(*ϕ*) determines whether a pathogen with a low spillover rate or high spillover rate is more likely to host jump. Second, at high levels of spillover the information gained about *ϕ* is exactly balanced by the high number of future opportunities for a host jump to occur. As a result, there is little to no difference in risk between pathogens that spill over at moderate rates and those that spill over at infinitely high rates. Lastly, when we explicitly separate the rate of spillover from the duration of opportunities for spillover, we see that host jump risk is strongly influenced by the latter only.

## Discussion

Here we have derived a general framework to characterize the relationship between the spillover rate of a zoonotic disease and the probability of that pathogen successfully jumping into a novel host. A novelty of this framework is that it does not rely on complex demographic models, but instead uses an extremely simple model that leverages data from past spillover events to make predictions about the outcome of future spillover events. We derive an analytical solution for this model when our prior distribution on *ϕ* (the probability that a given spillover event will result in a host jump) follows a beta distribution or mixture of beta distributions. The analytical solution to this model (Eq. 4) demonstrates that in addition to the prior, two composite factors shape the risk of a host jump; 1) the rate of spillover in the past *λ* times the duration of past opportunities for spillover *t*_*P*_, and 2) the rate of spillover in the future *cλ* times the time window over which future host jump risk is being assessed *t*_*F*_. The model shows that conditional on a pathogen having not yet jumped hosts, the risk of a future host jump declines with the first composite factor and increases with the second composite factor (Fig. 2D-F). We then use this model to ask which is the bigger threat, a pathogen that has rarely spilled over in the past and will rarely spill over in the future, or a pathogen that has frequently spilled over in the past and will frequently spill over in the future. We show that the answer to this question depends on our prior distribution on *ϕ*, and that when the rate of spillover is moderate or higher, the difference in spillover rate between pathogens becomes negligible. This is because while pathogens with high rates of spillover will have many future opportunities to host jump, there is also an increasing amount of evidence that *ϕ* is small, given that a host jump has not yet occurred. Lastly, we show that in contrast to spillover rate, the risk of a host jump is highly sensitive to the duration of past opportunities for spillover *t*_*P*_, with the risk rapidly increasing as pathogens become more novel (i.e. the opportunity for spillover arose recently). We thus conclude that novelty, but not spillover rate, is a strong predictor of host jump risk.

A key point revealed by our model is that host jump risk is highly sensitive to the duration of time in the past over which there were opportunities for spillover *t*_*P*_(Fig. 2J-L). The intuitive interpretation of this result is that pathogens that have not host jumped despite having a long opportunity to do so pose a lower host jump threat than pathogens that are just now gaining their first opportunities for spillover. In other words, novel zoonotic threats are riskier than long-established zoonotic threats. An important point, of course, is that pathogen novelty can arise in multiple ways due to, for example, newly created interactions between native hosts and novel hosts (18, 29), the emergence of a pathogen in an intermediate host (30), or a pathogen evolving suffciently far over time in its native host that it is now effectively a different threat than the version of it that was around in the past (9, 31).

Previous studies addressing host jump risk have largely revolved around identifying underlying drivers of host jump risk based on correlative approaches (21, 32–35). Here we instead utilize tools from Bayesian statistics to infer how host jump risk changes as a function of the rate of spillover. This approach allows us to explicitly connect a pathogen’s spillover rate with its probability that any given spillover event will result in a host jump (i.e., our parameter *ϕ*). This connection demonstrates that pathogens that have had higher rates of spillover in the past, particularly those that have been around for a while, must on average have lower values of *ϕ* because if that were not the case, they would have already completed a host jump. Consequently, our model reveals that spillover rate is a poor predictor of host jump risk.

Our finding that spillover rate is not predictive of host jump risk is driven in part by the competing impacts of spillover rate on host jump risk; higher rates of spillover lead to more opportunities for a host jump to occur, but also indicate that any given spillover is unlikely to result in a host jump (i.e. *ϕ* is small). This is because pathogens with high inherent rates of spillover must have small values of *ϕ* on average, as they would have already host jumped otherwise (Fig. 1E). As shown in Eq. 5, when there is a linear correlation between the rate of spillover in the past and in the future, host jump risk converges on a fixed constant strictly between zero and one as spillover rate gets large. Thus pathogens with moderate or high rates of spillover are nearly equally risky (see Supplementary text G for an analysis of how the prior on *ϕ* affects the speed of this convergence). In addition, as we show with Scenarios 1-3, our prior uncertainty in the value of *ϕ* determines whether we approach this fixed constant from above or from below (i.e. the riskiest pathogens can be those that spill over rarely or those that spill over frequently). Consequently, when we are unable to reliably quantify our uncertainty in *ϕ*, spillover is not useful in predicting host jump risk, and even when we are able to quantify its uncertainty, it is only moderately useful. These findings, however, do not dispute the conventional wisdom that spillover reduction decreases host jump risk. As Eq. 3 demonstrates, decreasing future spillover will decrease host jump risk regardless of how many times a pathogen has spilled over in the past (i.e. moving top to bottom in Fig. 2D-F). It is also worth noting that spillover reduction provides public health benefits even in the absence of host jump risk (24, 29).

Our model illustrates that there are scenarios where host jump risk can be maximized at high, low, or intermediate values of spillover (Scenarios 1, 2, and 3 respectively). Moreover, the analytical solution to our model (Eq. 4) demonstrates that host jump risk prediction is dependent on the shape parameters *a* and *b* that generate our prior on *ϕ* (i.e. our uncertainty in the probability that a given spillover event will result in a host jump). Therefore, if one were interested in adapting our model to quantify risk for a particular zoonotic pathogen of interest, the prior on *ϕ* could be tailored for that particular pathogen or group of pathogens. This approach could account for pathogen-specific details that may not be generalizable to all zoonotic pathogens, such as phylogenetic relationships (14, 35) and genomic markers for the ability to infect and spread in novel hosts (31, 36). Similarly, analyses of past instances of successful host jumps, (13, 33, 37), ranking risk factors (34, 38), and the basic reproductive number, *R*_0_ (32, 39) could be incorporated into these modeling efforts.

Our model results depend on three basic assumptions, namely: 1) the outcome of all spillover events are independent of one another, 2) the average probability that a spillover event results in a host jump (*ϕ*) is constant over time, and 3) the rates of spillover in the past and future are linearly correlated. Each of these assumptions could be violated in certain situations. Our first assumption could be violated if co-occurring spillover events would affect the probability of either spillover resulting in a host jump. This effect would disproportionately impact pathogens with high rates of spillover, presumably decreasing the likelihood of a host jump relative to our predictions. Our second assumption, that *ϕ* is constant over time, could be violated for example due to changes in population density, geographic connectivity, behavior, host immunity, or evolutionary pressure (40). If *ϕ* were to increase over time, the outcomes of past spillover events would give less information about the risk posed by future spillover events, and we therefore suspect that pathogens with high rates of spillover may pose a relatively greater threat than those with low rates of spillover, but this conclusion would likely depend on the precise way that *ϕ* is changing over time. Our third assumption that past and future spillover rates are linearly correlated seems likely to hold for most pathogens in general. If this assumption were violated, however, the probability of a host jump would no longer converge to a value between zero and one (dashed gold line in Fig.2G-I), and would instead converge to 0 or 1, depending on whether the correlation was sub-linear or greater than linear (see Supplementary text D).

Mechanistic models have been used extensively for predicting timing and location of spillover (17, 41–43), but this approach has not been used to the same extent for host jump risk prediction (34, 35).Here, we have proposed a model framework that we consider to be mechanistic in the sense that the outcome of the model arises directly from the processes contained in our model. However, incorporating more biological details could serve to further improve this approach in the future (10, 21, 32, 33, 44). For instance, features such as phylogenetic or genomic data (14, 30, 36, 45), host-pathogen characteristics (7, 46, 47), and evolutionary theory (32, 39, 48, 49) could all be integrated into characterizing our prior uncertainty of *ϕ*. Implementing such mechanisms could provide distinct advantages by accounting for biological processes and may be better equipped to preemptively identify zoonotic threats. Combining such a model with efforts to develop therapeutics to “prototype pathogens” could enable rapid pharmaceutical responses to spillover events that are likely to result in a host jump, which could play a significant role in future pandemic prevention (50).

Our model suggests that the relationship between spillover rates and host jump risk can be counter-intuitive: the pathogen that spills over most is not necessarily the greatest host jump threat, nor is it the case that exceptionally rare or undiscovered pathogens pose the greatest risk. Rather, the pathogens that pose the greatest risk of a host jump are those that are novel, meaning that opportunities for spillover only became possible recently. Rather than focusing on spillover rate, our model shows that incorporating system-specific factors regarding hosts-pathogen-environment interactions and evolutionary processes will therefore be necessary to more definitively characterize host jump risk for specific pathogens of interest. Efforts to mitigate future spillover will continue to be a powerful pandemic prevention tool (29), but our framework shows that spillover rate alone is a poor predictor of host jump risk.

## ACKNOWLEDGMENTS

D.K. and B.S. were supported by Institute of General Medical Sciences, National Institutes of Health (R01GM140459) and the UK Biotechnology and Biological Sciences Research Council as part of the joint NSF-NIH-USDA Ecology and Evolution of Infectious Diseases program, and by National Science Foundation grant DEB-2211322. The funders had no role in study design, data collection and analysis, decision to publish, or preparation of this article.

## Data and materials availability

All code used to produce and analyze these results is publicly available on a GitHub repository at https://github.com/43simony/HostJumpmodelCode

## Supporting Information Appendix (SI)

### I. Supplemental Information text

A. **Accounting for uniqueness of individual spillover events.**
B. **Poisson model derivation.**
C. **Count model derivation.**
D. **Analytical solution to the limit (Poisson model).**
1. ***Analytical solution to the limit for a beta mixture distribution (Poisson model).***
E. **Analytical solution to the limit (Count model).**
1. ***Analytical solution to the limit for a beta mixture distribution (Count model).***
F. **Supplementary Results.**
G. **Evaluating reasonable prior hyper-parameters.**
1. ***Results under changing prior hyper-parameters.***

