## Supplementary text for "Spillover rate is not a predictor of host jump risk, but spillover novelty is"

### II. Supplementary Text

**A. Accounting for uniqueness of individual spillover events.** In the main text, we presented Eq.1, which describes the probability that no host jumps occur in some length of time with a fixed rate of spillover $\lambda$. In that equation, we assumed that every spillover event had the same probability $\phi$ of resulting in a host jump. Here we show that relaxing the assumption that all spillover events have the same probability of resulting in a host jump has no impact on our model.

We define $\phi_j$ as the probability that some spillover event $j$ results in a successful host jump. The value of a given $\phi_j$ depends on host-pathogen-environment interactions as well as temporal and individual level variation that might affect the likelihood of a host jump. For example, individuals may differ in their probability of becoming infected or in their transmissibility, so the particular individuals that become infected may change the probability that the given spillover event results in a host jump. Not all values of $\phi_j$ are equally likely, so a given value of $\phi_j$ can be theoretically characterized as a random draw from some distribution $\Phi$ with density function $f_\Phi(\phi_j)$. Therefore, the probability that a single spillover event ($N = 1$) results in a successful host jump ($H > 0$, or more specifically here $H = 1$) can be quantified as the product of $\phi_j$ and the probability density function for this particular value of $\phi_j$, integrated over all possible values of $\phi$.

$$P(H > 0|N = 1, \Phi) = \int_0^1 \phi_j \cdot f_\Phi(\phi_j) d\phi_j \quad \text{[S1]}$$

$$= E(\Phi) = \phi \quad \text{[S2]}$$

We see that this is the definition of the expected value (or mean) of a distribution, and we define $\phi$ as the mean of $\Phi$. If we expand this to any number of spillover events, the probability that $N$ spillover events fail to result in a host jump ($H = 0$) where the vector of probabilities that the $j^{th}$ spillover event results in a host jump ($\vec{\phi}_j$) can be written as:

$$P(H = 0|N = 1, \vec{\phi}_j) = \prod_{j=1}^{N} (1 - P(H > 0|N = 1, \phi_j)) \quad \text{[S3]}$$

$$= \prod_{j=1}^{N} (1 - \phi_j) \quad \text{[S4]}$$

Due to the stochastic nature of spillover and the ensuing disease dynamics, it is effectively impossible to know the true $\phi_j$ for a given spillover event, so we instead multiply Eq.S4 by the probability that $\phi_j$ takes a particular value and integrate over all values of $\phi_j$. Because we assume that each spillover event is independent, we can apply this to each instance of spillover individually such that

$$P(H = 0|N, \Phi) = \prod_{j=1}^{N} \int_0^1 (1 - \phi_j) \cdot f_\Phi(\phi_j) d\phi_j \quad \text{[S5]}$$

$$= \prod_{j=1}^{N} \int_0^1 f_\Phi(\phi_j) d\phi_j - \int_0^1 \phi_j \cdot f_\Phi(\phi_j) d\phi_j \quad \text{[S6]}$$

Using the definition of $\phi$ from Eq.S2 and the fact that $f_\Phi(\phi_j)$ is a proper density function and therefore integrates to 1 over its support, we get

$$P(H = 0|N, \Phi) = \prod_{j=1}^{N} 1 - \phi \quad \text{[S7]}$$

$$P(H = 0|N, \Phi) = (1 - \phi)^N \quad \text{[S8]}$$

We see that this equation exactly describes the binomial probability of observing no host jumps in some number of spillover events ($N$), if we assume that the mean of all possible values of $\phi_j$ does not change over time. Since this equation holds for all positive integer values of $N$, we can then use the law of total probability to expand this equation so that the number of spillover events in a given amount of time is stochastic. If we explicitly assume that the number of spillover events in some time interval ($t$) follows a Poisson process where $\lambda$ is the pathogen's rate of spillover, we arrive at equation 1, and we show this derivation in Supplementary text B. Therefore, we may equivalently express the probability of no host jumps ($H = 0$) for a pathogen with spillover rate $\lambda$ in some time interval $t$ when each spillover event has a unique probability of resulting in a host jump $\phi_j$ as a function of the mean of all possible values of $\phi_j$, which we refer to as $\phi$.

**B. Poisson model derivation.** In the main text, we presented a model for the probability of a successful host jump in some future time horizon ($t_F$) that depends on a pathogen's inherent rate of spillover ($\lambda$), the duration of the shared ecological history between the novel host and the pathogen ($t_P$), and the probability that an individual spillover event results in a successful host jump ($\phi$). Here, we present a full generalized derivation of this model and provide an analytical solution for the case where the prior on $\phi$ follows a beta or mixture of beta distributions.

In what follows, we will derive a model for the probability of at least one successful host jump ($H_F > 0$) occurring in a fixed future time horizon ($t_P$) as a function of a pathogen's rate of spillover ($\lambda$), the length of time that spillover was possible in the past ($t_P$), the number of past host jumps ($H_P$), and our uncertainty in the probability that a spillover results in a successful host jump $\pi(\phi)$. We make the same assumptions as in the main text that the outcome of each spillover event is independent of all others, and that $\phi$ is not changing over time. Here, we assume that the number of spillover events in the past and future follow Poisson processes with rates $\lambda t_P$ and $c\lambda t_F$ respectively, so the number of spillovers in the past ($N$) and future ($M$) can be expressed as

$$N \sim Pois(\lambda t_P) \quad \text{[S9]}$$

$$M \sim Pois(c\lambda t_F) \quad \text{[S10]}$$

Equation 1 in the main text, which quantifies the probability that exactly $H_P$ host jumps occur, conditional on $N$ and $\phi$ can the be derived using the law of total probability such that

DRAFT

$$P(H_P|\lambda, t_P, \phi) = \sum_{N=H_P}^{\infty} P(H_P|\phi, N) \cdot P(N|\lambda, t_P) \quad \text{[S11]}$$

$$P(H_P|\lambda, t_P, \phi) = \frac{1}{C_{norm}} \sum_{N=H_P}^{\infty} \binom{N}{H_P} \phi^{H_P} (1-\phi)^{N-H_P} \cdot \frac{(\lambda t_P)^N e^{-\lambda t_P}}{N!} \quad \text{[S12]}$$

$$C_{norm} = 1 - \sum_{k=0}^{H_P - 1} \frac{(\lambda t_P)^k e^{-\lambda t_P}}{k!} \quad \text{[S13]}$$

where $C_{norm}$ is a normalizing constant, which is necessary due to the constraint that $N \geq H_P$, and is only necessary when $H_P > 0$. We can continue to simplify the infinite sum in Eq.S13 such that

$$P(H_P|\lambda, t_P, \phi) = \frac{e^{-\lambda t_P} (\phi \lambda t_P)^{H_P}}{C_{norm} \cdot H_P!} \sum_{N=H_P}^{\infty} \frac{(\lambda t_P - \phi \lambda t_P)^{N-H_P}}{(N-H_P)!} \quad \text{[S14]}$$

$$= \frac{e^{-\lambda t_P} (\phi \lambda t_P)^{H_P}}{C_{norm} \cdot H_P!} \sum_{N-H_P=0}^{\infty} \frac{(\lambda t_P - \phi \lambda t_P)^{N-H_P}}{(N-H_P)!} \quad \text{[S15]}$$

$$= \frac{e^{-\lambda t_P} (\phi \lambda t_P)^{H_P}}{C_{norm} \cdot H_P!} \cdot e^{\lambda t_P - \phi \lambda t_P} \quad \text{[S16]}$$

Following these simplifying steps, we use the fact that that the sum has the form $\sum_{n=0}^{\infty} \frac{x^n}{n!}$, which is the series definition of the exponential function. Using this property and simplifying this expression further, we get:

$$P(H_P|\lambda, t_P, \phi) = \frac{(\phi \lambda t_P)^{H_P}}{C_{norm} \cdot H_P!} \cdot e^{-\phi \lambda t_P} \quad \text{[S17]}$$

Equation S17 therefore gives a generalized form of Eq. 1 from the main text that allows for a non-zero number of past host jumps, given some rate of spillover ($\lambda t_P$) and the probability that a spillover results in a host jump ($\phi$). We can then use this equation to derive our model when the number of past and future spillover events are Poisson distributed with a fixed rate. Using Eq. S17 in Bayes' theorem and simplifying, the posterior distribution can be expressed as:

$$\pi(\phi|\lambda, t_P, H_P) = \frac{\pi(\phi) \cdot P(H_P|\lambda, t_P, \phi)}{\int_0^1 \pi(\phi) \cdot P(H_P|\lambda, t_P, \phi) d\phi} \quad \text{[S18]}$$

$$\pi(\phi|\lambda, t_P, H_P) = \frac{\pi(\phi) \cdot \frac{(\phi \lambda t_P)^{H_P}}{C_{norm} \cdot H_P!} e^{-\phi \lambda t_P}}{\int_0^1 \pi(\phi) \cdot \frac{(\phi \lambda t_P)^{H_P}}{C_{norm} \cdot H_P!} e^{-\phi \lambda t_P} d\phi} \quad \text{[S19]}$$

$$\pi(\phi|\lambda, t_P, H_P) = C \cdot \pi(\phi) \cdot e^{-\phi \lambda t_P} \phi^{H_P} \quad \text{[S20]}$$

In general, it is not possible to analytically evaluate the integral in the denominator for any particular prior, so numerical integration is often required to determine this normalizing constant. However, when the prior follows a beta distribution, this constant $C$ can be expressed in terms of the confluent hypergeometric function (28) and the posterior can be written as

$$\pi(\phi|\lambda, t_P, H_P) = \frac{\pi(\phi) \cdot e^{-\phi \lambda t_P} \phi^{H_P}}{\mathcal{M}(a + H_P, b + a + H_P, -\lambda t_P)} \quad \text{[S21]}$$

We may also substitute this form of the posterior in Eq. S20 and Eq. S17 to Eq. 3, so the full model can be expressed as:

$$P(H_F > 0|\pi(\phi), \lambda, t_P, c\lambda, t_F, H_P) = 1 - \int_0^1 \pi(\phi|\lambda, t_P, H_P) \cdot P(H_F = 0|c\lambda t_F, \phi) d\phi \quad \text{[S22]}$$

In the case that the prior follows a beta distribution, we can use the analytic form of the posterior distribution, and find that this expression can be simplified further using the confluent hypergeometric function such that

$$P(H_F > 0|\pi(\phi), \lambda, t_P, c\lambda, t_F, H_P) = 1 - \int_0^1 \frac{\pi(\phi) \cdot e^{-\phi \lambda t_P} \phi^{H_P}}{\mathcal{M}(a + H_P, b + a + H_P, -\lambda t_P)} \cdot e^{-\phi c \lambda t_F} d\phi \quad \text{[S23]}$$

$$= 1 - \frac{1}{\mathcal{M}(a + H_P, b + a + H_P, -\lambda t_P)} \int_0^1 \pi(\phi) \cdot e^{-\phi \lambda (t_P + c t_F)} \phi^{H_P} d\phi \quad \text{[S24]}$$

$$= 1 - \frac{\mathcal{M}(a + H_P, b + a + H_P, -\lambda (t_P + c t_F))}{\mathcal{M}(a + H_P, b + a + H_P, -\lambda t_P)} \quad \text{[S25]}$$

DRAFT

Additionally we may instead define $\pi(\phi)$ as a mixture of $K$ beta distributions such that

$$\pi(\phi) \sim \sum_{k=1}^{k} \omega_k \cdot \text{Beta}(a_k, b_k) \tag{S26}$$

$$\sum_{k=1}^{k} \omega_k = 1 \tag{S27}$$

Following a similar set of steps to those described for the single beta distribution, we derive

$$P(H_F > 0|\pi(\phi), \lambda, t_P, c\lambda, t_F, H_P) = 1 - \frac{\sum_{k=1}^{K} \omega_k C_k \mathcal{M}(a_k + H_P, b_k + a_k + H_P, -\lambda(t_P + t_F))}{\sum_{k=1}^{K} \omega_k C_k \mathcal{M}(a_k + H_P, b_k + a_k + H_P, -\lambda t_P)} \tag{S28}$$

$$C_k = \frac{\Gamma(a_k + H_P)\Gamma(a_k + b_k)}{\Gamma(a_k)\Gamma(b_k + a_k + H_P)} \tag{S29}$$

Therefore, we have derived a solution to the model when the number of past and future spillover events are defined by Poisson distributions with fixed rates. This addition to the model also allows for cases where there have been host jumps in the past, similar to the main model. In addition, we note that it is possible to mix model structures such that the past may have a fixed number of spillover events ($N$) while future spillover events depend on a fixed rate ($c\lambda t_F$) and vice versa. This plug-and-play feature could be useful in contexts when the uncertainty in the number of spillovers differs between the past and the future.

**C. Count model derivation.** In addition to our model which describes the probability of at least one future host jump based on past and future rates of spillover, we also consider a simpler form which instead relies on counts of spillover events. Here we derive an analytical solution for our model, which describes the probability of at least one host jump in some number of future spillover events ($M$). We condition this probability on the number of past spillovers ($N$), the number of past spillovers that resulted in a host jump ($H_P$), and our uncertainty in the probability that a single spillover results in a host jump $\pi(\phi)$. In the main text, we only consider cases where no past spillovers have resulted in a host jump ($H_P = 0$), but in our derivations here, we show that our model is also valid in cases where past host jumps have occured.

First, we define our prior using a beta distribution, such that $\pi(\phi) \sim \text{Beta}(a,\ b)$. We make the same assumptions as in the main text that the outcome of each spillover event is independent of all others, and that $\phi$ is not changing over time. Under these assumptions, we can use a binomial distribution to quantify the probability that exactly $H_P$ host jumps occured in $N$ past spillover events, and each spillover had some fixed probability $\phi$ of resulting in a host jump. This yields:

$$P(H_P|N, \phi) = \binom{N}{H_P} \phi^{H_P} (1 - \phi)^{N - H_P} \tag{S30}$$

Using Bayes' theorem, we can update $\pi(\phi)$ using the outcomes of these past spillover events, where the posterior distribution has the form

$$\pi(\phi|H_P, N) = \frac{\pi(\phi) \cdot P(H_P|N, \phi)}{\int_0^1 \pi(\phi) \cdot P(H_P|N, \phi) d\phi} \tag{S31}$$

Using the fact that the beta and binomial distributions have a conjugate relationship, it is known that the posterior distribution will follow a beta distribution, parameterized such that

$$\pi(\phi|H_P, N) \sim \text{Beta}(a + H_P,\ b + N - H_P) \tag{S32}$$

We then use the posterior distribution in Eq.S32 to compute the probability of at least one future host jump ($H_F > 0$) in some number of future spillover events ($M$). We express this instead as one minus the probability that no host jumps occur in $M$ spillover events. The probability of no host jumps in $M$ future spillovers can be written as the product of the posterior distribution from Eq.S32 and Eq.S30 with $M$ in place of $N$ and $H_F$ in place of $H_P$. We account for our posterior uncertainty on the value of $\phi$ by integrating over $[0, 1]$, which encompasses all possible values of $\phi$. This gives

$$P(H_F > 0|\pi(\phi), N, M, H_P) = 1 - P(H_F = 0|\pi(\phi), N, M, H_P) \tag{S33}$$

$$= 1 - \int_0^1 \pi(\phi|H_P, N) \cdot P(H_F = 0|M, \phi) d\phi \tag{S34}$$

If we substitute in the posterior and the right-hand side of Eq.S30, we see that the integrand is proportional to the PDF of a beta distribution, so the integral in S34 can be computed analytically, yielding:

$$P(H_F > 0|\pi(\phi), N, M, H_P) = 1 - \frac{\Gamma(b + N + M - H_P) \cdot \Gamma(a + b + N)}{\Gamma(a + b + N + M) \cdot \Gamma(b + N - H_P)} \tag{S35}$$

As before, we can instead define a prior distribution as a mixture of $K$ independent beta distributions where

$$\pi(\phi) \sim \sum_{k=1}^{k} \omega_k \cdot \text{Beta}(a_k, b_k) \tag{S36}$$

$$\sum_{k=1}^{k} \omega_k = 1 \tag{S37}$$

Again using the conjugate relationship between the beta and binomial distributions following the same sequence of steps as before, we derive

DRAFT

$$P(H_F > 0|\pi(\phi), N, M, H_P) = 1 - \sum_{k=1}^{K} \frac{\omega_k}{C(N)} \frac{\Gamma(a_k+b_k)}{\Gamma(a_k)\Gamma(b_k)} \frac{\Gamma(b_k+N+M-H_P)\cdot\Gamma(a_k+H_P)}{\Gamma(a_k+b_k+N+M)} \quad \text{[S38]}$$

$$C(N) = \sum_{k=1}^{K} \omega_k \frac{\Gamma(a_k+b_k)}{\Gamma(a_k)\Gamma(b_k)} \frac{\Gamma(a_k+H_P)\Gamma(b_k+N-H_P)}{\Gamma(a_k+b_k+N)} \quad \text{[S39]}$$

Therefore, we have closed form analytical solutions to our model when our prior on $\phi$ follows a beta distribution or mixture of betas, even in cases where past host jumps occured ($H_P > 0$). As a result, our model is not restricted to groups of pathogens that have not previously host jumped, and this logic could be similarly applied to cases in which public health responses might have prevented a successful host jump. More importantly, this allows us to evaluate host jump risk as a function of past spillovers, past host jumps, and future host jumps when $\phi$ follows a beta distribution without the need to rely on numerical integration or simulations. Our exact analytical solution allows us to avoid any computational errors in our solution that may arise in these other approaches.

**D. Analytical solution to the limit (Poisson model).** Here, we consider how increasing the rate of spillover relates to host jump risk, specifically when the rate of spillover approaches infinitely high values. Intuitively, one might expect that infinite spillover would always result in a host jump with probability 1 unless the probability of host jump given spillover was exactly 0. However, our main results suggest that host jump risk converges to a value between zero and one at high rates of spillover. To validate this observation, we compute the limit of our analytical solution as spillover rates become infinitely high.

Using the analytical solution to our model for the case with a fixed rate of spillover over some past and future time intervals, we can take the limit as $\lambda$ approaches infinity. As in the main text, we use $\lambda$ to represent the rate of past spillover and $f(\lambda)$ to represent the rate of future spillover. In the main text, we assumed a linear relationship such that $f(\lambda) = c\lambda$, but we relax this assumption here to demonstrate the potential effects of non-linear relationships as well. We can then express the probability of a host jump as the rate of spillover becomes infinitely high as

$$P_\infty(H_F > 0|\bullet) = \lim_{\lambda\to\infty} 1 - \frac{\mathcal{M}\left(a, a+b, -\lambda(f(\lambda)\cdot t_F + t_P)\right)}{\mathcal{M}\left(a, a+b, -\lambda t_P\right)} \quad \text{[S40]}$$

where $\bullet$ represents the model parameters, as in the main text. We can then use the asymptotic properties of the confluent hypergeometric function. Because all of our model parameters are positive real numbers, we can apply the asymptotic behavior of the confluent hypergeometric function as the parameter $|z|$ approaches infinity (28). Namely,

$$\mathcal{M}\left(a, b, z\right) \sim \frac{\Gamma(b)}{\Gamma(b-a)}(-z)^{-a} \quad \text{[S41]}$$

We can then express the limit in terms of this asymptotic form and simplify so

$$\lim_{\lambda\to\infty} 1 - \frac{\mathcal{M}\left(a, a+b, -(f(\lambda)\cdot t_F + \lambda t_P)\right)}{\mathcal{M}\left(a, a+b, -\lambda t_P\right)} = \lim_{\lambda\to\infty} 1 - \frac{\frac{\Gamma(a+b)}{\Gamma(b)}\left((f(\lambda)\cdot t_F + \lambda t_P)\right)^{-a}}{\frac{\Gamma(a+b)}{\Gamma(b)}(\lambda t_P)^{-a}} \quad \text{[S42]}$$

$$= \lim_{\lambda\to\infty} 1 - \frac{(\lambda t_P)^a}{(f(\lambda)\cdot t_F + \lambda t_P)^a} \quad \text{[S43]}$$

Using this property, we can express the probability of at least one host jump as the rate of spillover becomes infinitely large as

$$\begin{aligned} P_\infty(H_F > 0|\bullet) &= 1 - \lim_{\lambda\to\infty} \frac{(\lambda t_P)^a}{(f(\lambda)\cdot t_F + \lambda t_P)^a} \\ &= 1 - \lim_{\lambda\to\infty} \frac{1}{\left(\frac{f(\lambda)}{\lambda}\cdot\frac{t_F}{t_P} + 1\right)^a} \end{aligned}$$

Here, we notice that the functional relationship between past and future spillover rates determines wether the probability of a future host jump approaches zero, one, or an intermediate value. Namely, if $f(\lambda)$ grows at a rate that is greater than linear (i.e., dramatic increase of spillover rates in the future), the limit equals zero (and thus the probability of a future host jump equals one). In contrast, if $f(\lambda)$ grows at a rate that is less than linear (i.e., diminishing increases in future spillover rates), the limit equals one (so the probability of a future host jump is zero). However, under a linear relationship (i.e., $f(\lambda) = c\lambda$), this equation simplifies such that

$$\begin{aligned} P_\infty(H_F > 0|\bullet) &= 1 - \frac{1}{\left(c\frac{t_F}{t_P} + 1\right)^a} \lim_{\lambda\to\infty} \frac{\lambda^a}{\lambda^a} \\ &= 1 - \frac{1}{(cT+1)^a} \end{aligned}$$

where $c$ is the ratio between past and future spillover rates, $T = \frac{t_f}{t_P}$ is the ratio between the future time interval over which host jump risk is being evaluated and the duration in which a host jump could have occured in the past, and $a$ is the first shape parameter in the beta prior, $\pi(\phi)$.

***2. Analytical solution to the limit for a beta mixture distribution (Poisson model).*** We also consider the value of this limit when the prior follows a mixture of beta distributions. Using the analytical solution for a beta mixture prior derived in B when the number of host jumps

$H_P = 0$, this limit can be written for any mixture of $K$ beta distributions such that

$$P_\infty(H_F > 0|\bullet) = 1 - \lim_{\lambda\to\infty} \sum_{k=1}^{K} \frac{\omega_k}{C(\lambda)} \mathcal{M}(a, a+b, -(f(\lambda)\cdot t_F + \lambda t_P)) \quad \text{[S44]}$$

$$C(\lambda) = \sum_{j=1}^{K} \omega_j \mathcal{M}(a, a+b, -\lambda t_P) \quad \text{[S45]}$$

Where as before, we use $\lambda$ to represent the rate of past spillover, we define the rate of future spillover as $f(\lambda)$, and $\bullet$ represents the model parameters. In what follows, we will assume that $f(\lambda) = c\lambda$, as our previous derivations suggest nonlinear relationships would still result in convergence to a value of zero or one depending on the direction of the relationship. Rewriting Eq.S45 under this linear relationship gives

$$P_\infty(H_F > 0|\bullet) = 1 - \lim_{\lambda\to\infty} \sum_{k=1}^{K} \frac{\omega_k \mathcal{M}(a_k, a_k + b_k, -\lambda(ct_F + t_P))}{\sum_{j=1}^{K} \omega_j \mathcal{M}(a_j, a_j + b_j, -\lambda t_P)} \quad \text{[S46]}$$

Using the fact that the limit of a sum is equal to the sum of the limit of each individual term, we can rewrite this as

$$P_\infty(H_F > 0|\bullet) = 1 - \sum_{k=1}^{K} \lim_{\lambda\to\infty} \frac{\omega_k \mathcal{M}(a_k, a_k + b_k, -\lambda(ct_F + t_P))}{\sum_{j=1}^{K} \omega_j \mathcal{M}(a_j, a_j + b_j, -\lambda t_P)} \quad \text{[S47]}$$

$$= 1 - \sum_{k=1}^{K} \lim_{\lambda\to\infty} \frac{\omega_k T_k(\lambda(ct_F + t_P))}{\sum_{j=1}^{K} \omega_j D_j(\lambda t_P)} \quad \text{[S48]}$$

where $T_k(\lambda)$ and $D_j(\lambda)$ are shorthand representations of the terms in the numerator and denominator respectively. Because of the sum in the denominator, computing this limit is not straightforward, so we instead find an upper and lower bound for this limit. For a sufficiently large value of $\lambda$, we want to show that that there is some $j^*$ such that $D_{j^*}(\lambda) \geq D_j(\lambda)$ for all values of $j$. Representing this using limits, we have

$$\lim_{\lambda\to\infty} D_j(\lambda) \leq \lim_{\lambda\to\infty} D_{j^*}(\lambda) \quad \text{[S49]}$$

$$\lim_{\lambda\to\infty} \frac{D_j(\lambda)}{D_{j^*}(\lambda)} \leq 1 \quad \text{[S50]}$$

By substituting in the expression for $D_j(\lambda)$ and using the asymptotic relationship for the confluent hypergeometric function,

$$\mathcal{M}(a, b, z) \sim \frac{\Gamma(b)}{\Gamma(b-a)}(-z)^{-a} \quad \text{[S51]}$$

$$\mathcal{M}(a, a+b, z) \sim \frac{\Gamma(a+b)}{\Gamma(b)}(-z)^{-a} \quad \text{[S52]}$$

we evaluate the limit on the left-hand side of Eq.S50 to express this inequality in terms of the parameters of the beta mixture distribution.

$$\lim_{\lambda\to\infty} \frac{\omega_j \mathcal{M}(a_j, a_j + b_j, -\lambda)}{\omega_{j^*} \mathcal{M}(a_{j^*}, a_{j^*} + b_{j^*}, -\lambda)} \quad \text{[S53]}$$

$$= \frac{\omega_j \Gamma(b_j)\Gamma(a_{j^*} + b_{j^*})}{\omega_{j^*}\Gamma(b_{j^*})\Gamma(a_j + b_j)} \lim_{\lambda\to\infty} \frac{\lambda^{-a_j}}{\lambda^{-a_{j^*}}} \quad \text{[S54]}$$

$$= C \cdot \lim_{\lambda\to\infty} \lambda^{a_{j^*} - a_j} \quad \text{[S55]}$$

From this, we see that the limit does not exist when $a_{j^*} > a_j$, is equal to $C$ when $a_{j^*} = a_j$, and zero when $a_{j^*} < a_j$. We see that the inequality in Eq.S50 is always satisfied when $a_{j^*} < a_j$. Thus, for a significantly large $\lambda$, we can define the index of the largest value of $D_j(\lambda)$ such that $j^* = \{j \mid a_j = min(\hat{a}_j)\}$. Even when there is not a unique smallest value of $a_j$, any value in the set $j^*$ would provide almost equivalent upper bounds. To ensure the tightest possible upper and lower bounds it is necessary to find the unique value in $j^*$ that maximizes $D_j(\lambda)$. This can be determined a priori, as this value only depends on the beta mixture parameters, but we find $D_j(\lambda)$ will be maximized for the $j^*$ with the largest value of $b_{j^*}$. We then define this unique index $(\Omega)$ such that $D_\Omega \geq D_j$ for all values of $j$. This is reasonable as, decreasing the beta shape parameter $a$ and increasing the beta shape parameter $b$ both drive the mean of a beta distribution towards zero, which would have the greatest likelihood in a mixture of betas when the number of spillovers is large.

Using this observation, we define an upper and lower bound on the limit from Eq.S48 when $\lambda$ is sufficiently large, such that

$$\sum_{k=1}^{K} \lim_{\lambda\to\infty} \frac{\omega_k T_k(\lambda)}{D_\Omega(\lambda)} \leq \sum_{k=1}^{K} \lim_{\lambda\to\infty} \frac{\omega_k T_k(\lambda)}{\sum_{j=1}^{K} \omega_j D_j(\lambda)} \leq \sum_{k=1}^{K} \lim_{\lambda\to\infty} \frac{\omega_k T_k(\lambda)}{\omega_\Omega D_\Omega(\lambda)} \quad \text{[S56]}$$

For the upper bound term, we substitute the expressions for the numerator and denominator, so our upper bound limit can be expressed as

$$\sum_{k=1}^{K} \lim_{\lambda\to\infty} \frac{\omega_k \mathcal{M}(a_k, a_k + b_k, -\lambda(ct_F + t_P))}{\sum_{j=1}^{K} \omega_j \mathcal{M}(a_j, a_j + b_j, -\lambda t_P)} \leq \sum_{j=1}^{K} \lim_{\lambda\to\infty} \frac{\omega_k \mathcal{M}(a_k, a_k + b_k, -\lambda(ct_F + t_P))}{\omega_\Omega \mathcal{M}(a_\Omega, a_\Omega + b_\Omega, -\lambda t_P)} \quad \text{[S57]}$$

DRAFT

Using the asymptotic form of the confluent hypergeometric function and the fact that $a_\Omega \leq a_k$ for all values of $k$, we see that every limit term within the sum will be zero, except when $k \in j^*$, and applying these results, we can rewrite and compute the limit such that

$$\sum_{k=1}^{K} \lim_{\lambda\to\infty} \frac{\omega_k \mathcal{M}(a_k, a_k+b_k, -\lambda(ct_F+t_P))}{\sum_{j=1}^{K} \omega_j \mathcal{M}(a_j, a_j+b_j, -\lambda t_P)} \leq \left(\frac{t_P}{ct_F+t_P}\right)^{a_\Omega} \sum_{k\in j^*} \frac{\omega_k \Gamma(b_\Omega)\Gamma(a_k+b_k)}{\omega_\Omega \Gamma(b_k)\Gamma(a_\Omega+b_\Omega)} \quad \text{[S58]}$$

$$\leq \frac{1}{(cT+1)^{a_\Omega}} \sum_{k\in j^*} \frac{\omega_k \Gamma(b_\Omega)\Gamma(a_k+b_k)}{\omega_\Omega \Gamma(b_k)\Gamma(a_\Omega+b_\Omega)} \quad \text{[S59]}$$

The lower bound can be computed similarly, such that

$$\frac{1}{(cT+1)^{a_\Omega}} \sum_{k\in j^*} \frac{\omega_k \Gamma(a_k+b_k)\Gamma(b_\Omega)}{\Gamma(a_\Omega+b_\Omega)\Gamma(b_k)} \leq \sum_{k=1}^{K} \lim_{\lambda\to\infty} \frac{\omega_k \mathcal{M}(a_k, a_k+b_k, -\lambda(ct_F+t_P))}{\sum_{j=1}^{K} \omega_j \mathcal{M}(a_j, a_j+b_j, -\lambda t_P)} \quad \text{[S60]}$$

Thus, when there is a single smallest value of $a_j$, the lower and upper bounds are exactly equal. For this case, we have an exact analytical solution for the limit when the prior distribution follows a mixture of beta distributions, such that

$$P_\infty(H_F > 0|\bullet) = 1 - \frac{1}{(cT+1)^{a_\Omega}} \quad \text{[S61]}$$

When there is not a unique smallest value of $a_j$, the lower and upper bounds will never be exactly equal, so we can only analytically compute bounds for the limit. Since we are interested in the probability of at least one host jump in the future, we only consider the upper bound on this probability as a conservative approach. To do this, we substitute our analytically computed lower bound, which gives

$$P_\infty(H_F > 0|\bullet) = 1 - \sum_{k=1}^{K} \lim_{\lambda\to\infty} \frac{\omega_k \mathcal{M}(a_k, a_k+b_k, -\lambda(ct_F+t_P))}{\sum_{j=1}^{K} \omega_j \mathcal{M}(a_j, a_j+b_j, -\lambda t_P)} \quad \text{[S62]}$$

$$\leq 1 - \frac{1}{(cT+1)^{a_\Omega}} \sum_{k\in j^*} \frac{\omega_k \Gamma(a_k+b_k)\Gamma(b_\Omega)}{\Gamma(a_\Omega+b_\Omega)\Gamma(b_k)} \quad \text{[S63]}$$

The gamma functions in this expression are strictly positive because parameters $\omega_k$, $a_k$, $b_k$, and $c$ are all greater than zero for all values of $k$, every term, we notice

$$\frac{1}{(cT+1)^{a_\Omega}} \sum_{k\in j^*} \frac{\omega_k \Gamma(a_k+b_k)\Gamma(b_\Omega)}{\Gamma(a_\Omega+b_\Omega)\Gamma(b_k)} > 0 \quad \text{[S64]}$$

Therefore, we have demonstrated that, when there is a linear correlation between the number of spillover events in the past and future, the probability of a host jump saturates to a value between zero and one as the rate of spillover approaches infinite values, and this is valid for any mixture of beta distributions. We find that this value depends heavily on the minimum value of the shape parameter $a_j$ in the mixture of beta distributions, although the other parameters are also relevant to the computed upper bound when the smallest $a_j$ is not unique.

**E. Analytical solution to the limit (Count model).** Here, we consider how increasing the rate of spillover relates to host jump risk, specifically when the rate of spillover approaches infinitely high values. Intuitively, one might expect that infinite spillover would always result in a host jump with probability 1 unless the probability of host jump given spillover was exactly 0. However, our main results suggest that host jump risk converges to a value between zero and one at high levels of spillover. To validate this observation, we compute the limit of our analytical solution as spillover rates become infinitely high.

Because this model has an analytical solution, we aim to quantify the long-run behavior of the curve when the number of past and future spillover events are linearly related. We define the number of future spillover events ($M$) as a linear function of past spillover events ($N$) such that $M = f(N)$ where $f(N)$ can be any differentiable function. Here, we focus on a linear relationship ($f(N) = cN$), but also consider how non-linear relationships would affect the value of this limit. Substituting this into our analytical solution and taking the limit as $N$ approaches infinity, we get:

$$\lim_{N\to\infty} \left(1 - \frac{\Gamma(b+N+M)\cdot\Gamma(a+b+N)}{\Gamma(a+b+N+M)\cdot\Gamma(b+N)}\right) \quad \text{[S65]}$$

$$= 1 - \lim_{N\to\infty} \left(\frac{\Gamma(b+f(N)+N)\cdot\Gamma(a+b+N)}{\Gamma(a+b+f(N)+N)\cdot\Gamma(b+N)}\right) \quad \text{[S66]}$$

In the trivial case when parameters $a$, $b$, $c$ are all positive, non-zero integers, we can rewrite this expression using the relationship between the gamma function and factorial expressions, where $(N-1)! = \Gamma(N)$. Using this relationship and simplifying the argument of the limit in Eq.S66:

$$\frac{\Gamma(b+(c+1)N)\cdot\Gamma(a+b+N)}{\Gamma(a+b+(c+1)N)\cdot\Gamma(b+N)} = \frac{(b+(c+1)N-1)!\cdot(a+b+N-1)!}{(a+b+(c+1)N-1)!\cdot(b+N-1)!} \quad \text{[S67]}$$

$$= \frac{(b+(c+1)N-1)!}{(a+b+(c+1)N-1)!} \frac{(a+b+N-1)!}{(b+N-1)!} \quad \text{[S68]}$$

$$= \frac{\prod_{k=1}^{a} a+b+N-k}{\prod_{k=1}^{a} a+b+(c+1)N-k} \quad \text{[S69]}$$

$$= \prod_{k=1}^{a} \frac{a+b+N-k}{a+b+(c+1)N-k} \quad \text{[S70]}$$

DRAFT

Substituting this simplified form back into Eq. S66,

$$1 - \lim_{N\to\infty} \frac{\Gamma(b+(c+1)N)\cdot\Gamma(a+b+N)}{\Gamma(a+b+(c+1)N)\cdot\Gamma(b+N)} = 1 - \lim_{N\to\infty} \prod_{k=1}^{a} \frac{a+b+N-k}{a+b+(c+1)N-k} \quad \text{[S71]}$$

Using properties of limits, the limit of a product is equal to the product of the limit for each term individually, given that each limit exists. Applying this property, we see that the limits of each term in the product do exist. Thus we can evaluate the limit and simplify, yielding

$$1 - \lim_{N\to\infty} \prod_{k=1}^{a} \frac{a+b+N-k}{a+b+(c+1)N-k} = 1 - \prod_{k=1}^{a} \lim_{N\to\infty} \frac{N+a+b-k}{(c+1)N+a+b-k} \quad \text{[S72]}$$

$$= 1 - \prod_{k=1}^{a} \frac{1}{(c+1)} \quad \text{[S73]}$$

$$= 1 - \left(\frac{1}{(c+1)}\right)^{a} \quad \text{[S74]}$$

Indeed, the probability of a host jump approaches a value between zero and one. However, this proof is only valid if parameters $a$, $b$, $c$ are all restricted to positive non-zero integer values, and we would like to prove that this claim holds for all valid parameter values. Since there are no straightforward simplifying steps when $a$, $b$, and $c$ are not integers, we use Stirling's approximation for the gamma function, which converges asymptotically to the gamma function as N approaches infinity. For now, we focus just on the limit term from Eq.S66, and applying Stirling's approximation yields:

$$\lim_{N\to\infty} \frac{\sqrt{2\pi(b+f(N)+N-1)}\left(\frac{b+f(N)+N-1}{e}\right)^{b+f(N)+N-1}\sqrt{2\pi(a+b+N-1)}\left(\frac{a+b+N-1}{e}\right)^{a+b+N-1}}{\sqrt{2\pi(a+b+f(N)+N-1)}\left(\frac{a+b+f(N)+N-1}{e}\right)^{a+b+f(N)+N-1}\sqrt{2\pi(b+N-1)}\left(\frac{b+N-1}{e}\right)^{b+N-1}} \quad \text{[S75]}$$

$$A = b+f(N)+N-1 \quad \text{[S76]}$$

$$B = a+b+N-1 \quad \text{[S77]}$$

$$C = a+b+f(N)+N-1 \quad \text{[S78]}$$

$$D = b+N-1 \quad \text{[S79]}$$

$$\lim_{N\to\infty} \sqrt{\frac{A\cdot B}{C\cdot D}} \cdot \frac{e^{C+D}}{e^{A+B}} \cdot \left(\frac{A^A B^B}{C^C D^D}\right) \quad \text{[S80]}$$

If we assume that the limit exists for each of these three terms, this limit can be expressed as the product of the limit of each term individually. We compute the limit for the first term in Eq.S80 for any functional relationship between past and future spillover such that

$$\mathcal{L}_1 = \lim_{N\to\infty} \sqrt{\frac{A\cdot B}{C\cdot D}} \quad \text{[S81]}$$

$$= \lim_{N\to\infty} \sqrt{\frac{(b+f(N)+N-1)\cdot(a+b+N-1)}{(a+b+f(N)+N-1)\cdot(b+N-1)}} \quad \text{[S82]}$$

$$= \lim_{N\to\infty} \sqrt{\frac{f(N)\cdot N+N^2+l.o.t.}{f(N)\cdot N+N^2+l.o.t.}} \quad \text{[S83]}$$

$$\mathcal{L}_1 = 1 \quad \text{[S84]}$$

Regardless the functional relationship $f(N)$, both the numerator and denominator will grow at the same rate as $N$ approaches infinity, and this limit will always approach a constant value of one. Since we have shown that $\mathcal{L}_1$ exists, we then compute the limit of the second term of Eq.S80.

$$\mathcal{L}_2 = \lim_{N\to\infty} \frac{e^{C+D}}{e^{A+B}} \quad \text{[S85]}$$

$$= \lim_{N\to\infty} \frac{e^{a+b+f(N)+N+b+N-2}}{e^{b+f(N)+N+a+b+N-2}} \quad \text{[S86]}$$

$$= \lim_{N\to\infty} \frac{e^{f(N)+2N+a+b+b-2}}{e^{f(N)+2N+b+a+b-2}} \quad \text{[S87]}$$

$$\mathcal{L}_2 = 1 \quad \text{[S88]}$$

DRAFT

We see that the numerator and denominator will always be exactly equal for any choice of $f(N)$, and this limit will also always evaluate to one. Again, because we have shown that $\mathcal{L}_2$ exists, we then compute the limit of the third term of Eq.S80.

$$\mathcal{L}_3 = \lim_{N\to\infty}\left(\frac{A^A B^B}{C^C D^D}\right) \tag{S89}$$

$$= \lim_{N\to\infty}\left(\frac{(b+f(N)+N-1)^{(b+f(N)+N-1)}(a+b+N-1)^{(a+b+N-1)}}{(a+b+f(N)+N-1)^{(a+b+f(N)+N-1)}(b+N-1)^{b+N-1}}\right) \tag{S90}$$

$$= \lim_{N\to\infty}\left(\frac{(b+f(N)+N-1)^{(b+f(N)+N-1)}(a+b+N-1)^{(a+b+N-1)}}{(a+b+f(N)+N-1)^{(a+b+f(N)+N-1)}(b+N-1)^{b+N-1}}\right)\cdot\frac{(b+N-1)^a}{(b+N-1)^a}\cdot\frac{(b+f(N)+N-1)^a}{(b+f(N)+N-1)^a} \tag{S91}$$

$$= \lim_{N\to\infty}\left(\frac{b+f(N)+N-1}{a+b+f(N)+N-1}\right)^{a+b+f(N)+N-1}\cdot\left(\frac{a+b+N-1}{b+N-1}\right)^{a+b+N-1}\cdot\left(\frac{b+N-1}{b+f(N)+N-1}\right)^{a} \tag{S92}$$

We simplify this expression to a product of three terms. Again, we assume the limit exists for each of the terms in Eq.S92 and evaluate each of these limits individually. For the first term, we get

$$l_1 = \lim_{N\to\infty}\left(\frac{b+f(N)+N-1}{a+b+f(N)+N-1}\right)^{a+b+f(N)+N-1} \tag{S93}$$

$$ln(l_1) = \lim_{N\to\infty}(a+b+f(N)+N-1)\cdot ln\left(\frac{b+f(N)+N-1}{a+b+f(N)+N-1}\right) \tag{S94}$$

$$= \lim_{N\to\infty}\frac{ln\left(\frac{b+f(N)+N-1}{a+b+f(N)+N-1}\right)}{\frac{1}{(a+b+f(N)+N-1)}} = \frac{0}{0} \tag{S95}$$

applying L'Hopitals rule and taking the derivatives of the numerator and denominator:

$$= \lim_{N\to\infty}\frac{\frac{f'(N)+1}{b+f(N)+N-1}-\frac{f'(N)+1}{a+b+f(N)+N-1}}{\frac{-(f'(N)+1)}{(a+b+f(N)+N-1)^2}}$$

$$= \lim_{N\to\infty}\frac{\frac{1}{b+f(N)+N-1}-\frac{1}{a+b+f(N)+N-1}}{\frac{-1}{(a+b+f(N)+N-1)^2}} \tag{S96}$$

$$= \lim_{N\to\infty}\frac{a}{(a+b+f(N)+N-1)(b+f(N)+N-1)}\frac{(a+b+f(N)+N-1)^2}{-1} \tag{S97}$$

$$= -a\lim_{N\to\infty}\frac{a+b+f(N)+N-1}{(b+f(N)+N-1)} = -a \tag{S98}$$

$$ln(l_1) = -a \tag{S99}$$

$$l_1 = e^{-a} \tag{S100}$$

Because we use L'Hopital's rule to evaluate this limit, we have an additional constraint on $f(N)$, which must be differentiable. If this condition is met, however, we see again that the chosen functional relationship between past and future spillover does not change the value of this limit. Again, this is because the chosen functional form appears in the numerator and denominator of the limit argument, so the rates of growth would exactly balance. Since $l_1$ exists, we use a similar sequence of steps to evaluate the limit of the second term in S92 such that

$$l_2 = \lim_{N\to\infty}\left(\frac{a+b+N-1}{b+N-1}\right)^{a+b+N-1} \tag{S101}$$

$$ln(l_2) = \lim_{N\to\infty}(a+b+N-1)\cdot ln\left(\frac{a+b+N-1}{b+N-1}\right) \tag{S102}$$

$$= \lim_{N\to\infty}\frac{ln\left(\frac{a+b+N-1}{b+N-1}\right)}{\frac{1}{(a+b+N-1)}} = \frac{0}{0} \tag{S103}$$

Applying L'Hopitals rule and taking the derivatives of the numerator and denominator:

$$= \lim_{N\to\infty}\frac{\frac{1}{b+N-1}-\frac{1}{a+b+N-1}}{\frac{-1}{(a+b+N-1)^2}}$$

$$= \lim_{N\to\infty}\frac{-a}{(a+b+N-1)(b+N-1)}\frac{(a+b+N-1)^2}{-1} \tag{S104}$$

$$= a\lim_{N\to\infty}\frac{a+b+N-1}{(b+N-1)} = a \tag{S105}$$

$$ln(l_2) = a \tag{S106}$$

$$l_2 = e^{a} \tag{S107}$$

DRAFT

The functional relationship between past and future spillover $f(N)$ is not present in this term, so alternative functional relationships would not affect the value of this limit. Since $l_2$ exists, we use a similar sequence of steps to evaluate the limit of the third term in S92 such that

$$l_3 = \lim_{N\to\infty}\left(\frac{b+N-1}{b+f(N)+N-1}\right)^a \quad [S108]$$

$$= \left(\lim_{N\to\infty}\frac{b+N-1}{b+f(N)+N-1}\right)^a \quad [S109]$$

Here, we see that the functional relationship between past and future spillover would affect the value of this limit. When the function $f(N)$ grows at a rate that is greater than linear, the limit evaluates to zero, and if $f(N)$ grows at a rate that is less than linear, the limit evaluates to one. We are interested in the case when the relationship between the number of past and future spillover events is exactly linear (i.e., $f(N) = cN$). If we substitute this expression into Eq.S109, we can evaluate the limit such that

$$l_3 = \lim_{N\to\infty}\left(\frac{b+N-1}{b+(c+1)N-1}\right)^a \quad [S110]$$

$$l_3 = \left(\frac{1}{(c+1)}\right)^a \quad [S111]$$

In any case, this limit does exist, so we can express the value of $\mathcal{L}_3$, noting that $l_1$ and $l_2$ exactly cancel one another such that

$$\mathcal{L}_3 = l_1 \cdot l_2 \cdot l_3 \quad [S112]$$

$$\mathcal{L}_3 = e^a e^{-a} \cdot l_3 \quad [S113]$$

$$\mathcal{L}_3 = l_3 \quad [S114]$$

Therefore, when the number of past and future spillover events are linearly correlated, we can express $\mathcal{L}_3$ as

$$\mathcal{L}_3 = \frac{1}{(c+1)^a} \quad [S115]$$

Because $\mathcal{L}_1$, $\mathcal{L}_2$, and $\mathcal{L}_3$ all exist, and $\mathcal{L}_1 = \mathcal{L}_2 = 1$ for any differentiable relationship $(f(N))$, we can express the value of the limit in Eq.S66 under a linear relationship as:

$$P_\infty(H_F > 0|\pi(\phi)) = 1 - \lim_{N\to\infty}\left(\frac{\Gamma(b+(c+1)\cdot N)\cdot\Gamma(a+b+N)}{\Gamma(a+b+(c+1)\cdot N)\cdot\Gamma(b+N)}\right) \quad [S116]$$

$$= 1 - \mathcal{L}_1 \cdot \mathcal{L}_2 \cdot \mathcal{L}_3 \quad [S117]$$

$$= 1 - l_3 \quad [S118]$$

$$= 1 - \frac{1}{(c+1)^a} \quad [S119]$$

From this, we can also see that this limit will take a value of either zero or one when $f(N)$ is non-linear . However, when this relationship is linear, we see that this expression is identical to the value derived in the case in Eq.S74 when the parameters $a$, $b$, $c$ were restricted to positive integer values. Because Stirling's approximation converges asymptotically to the gamma function for large values, we are able to exactly compute this limit and find a general form that is also valid for non-integer parameter values.

This derivation therefore demonstrates that the probability of a host jump saturates to a value between zero and one, even when the rate of spillover approaches infinite values. This result arises because a large number of past spillover events without a host jump drives the posterior density of $\phi$ towards zero, while a large number of future spillover events yields many opportunities for a future host jump. These competing factors exactly counterbalance one another in the limit when there is a linear correlation between the rate of spillover in the past and future. However, non-linear relationships between past and future spillover do not maintain this balance and do indeed converge exactly to zero or one.

***2. Analytical solution to the limit for a beta mixture distribution (Count model).*** We also consider the value of this limit when the prior follows a mixture of beta distributions. Using the analytical solution derived in Supplemental text C for $H_P = 0$, this limit can be written for any mixture of $K$ beta distributions such that

$$P_\infty(H_F > 0|\pi(\phi)) = 1 - \lim_{N\to\infty}\sum_{k=1}^{K}\frac{\omega_k}{C(N)}\frac{\Gamma(a_k+b_k)}{\Gamma(a_k)\Gamma(b_k)}\frac{\Gamma(b_k+N+M)\cdot\Gamma(a_k)}{\Gamma(a_k+b_k+N+M)} \quad [S120]$$

$$C(N) = \sum_{j=1}^{K}\omega_j\frac{\Gamma(a_j+b_j)}{\Gamma(a_j)\Gamma(b_j)}\frac{\Gamma(a_j)\Gamma(b_j+N)}{\Gamma(a_j+b_j+N)} \quad [S121]$$

To condense this equation, we use the beta function, which is defined as

$$B(a,b) = \frac{\Gamma(a)\Gamma(b)}{\Gamma(a+b)};\quad B^{-1}(a,b) = \frac{\Gamma(a+b)}{\Gamma(a)\Gamma(b)} \quad [S122]$$

Rewriting Eq.S121 with the beta function and substituting $M = cN$, we get

$$P_\infty(H_F > 0|\pi(\phi)) = 1 - \lim_{N\to\infty}\sum_{k=1}^{K}\frac{\omega_k B^{-1}(a_k,b_k)B(a_k,B_k+(c+1)N)}{\sum_{j=1}^{K}\omega_j B^{-1}(a_j,b_j)B(a_j,b_j+N)} \quad [S123]$$

DRAFT

Using the fact that the limit of a sum is equal to the sum of the limit of each individual term, we can rewrite this as

$$P_\infty(H_F > 0|\pi(\phi)) = 1 - \sum_{k=1}^{K} \lim_{N\to\infty} \frac{\omega_k B^{-1}(a_k, b_k) B(a_k, B_k + N + M)}{\sum_{j=1}^{K} \omega_j B^{-1}(a_j, b_j) B(a_j, b_j + N)} \quad [S124]$$

$$= 1 - \sum_{k=1}^{K} \lim_{N\to\infty} \frac{T_k(N)}{\sum_{j=1}^{K} D_j(N)} \quad [S125]$$

where $T_k(N)$ and $D_j(N)$ are shorthand representations of the terms in the numerator and denominator respectively. Because of the sum in the denominator, computing this limit is not straightforward, so we instead find an upper and lower bound for this limit. For a sufficiently large value of $N$, we want to show that that there is some $j^*$ such that $D_{j^*}(N) \geq D_j(N)$ for all values of $j$. Representing this using limits, we have

$$\lim_{N\to\infty} D_j(N) \leq \lim_{N\to\infty} D_{j^*}(N) \quad [S126]$$

$$\lim_{N\to\infty} \frac{D_j(N)}{D_{j^*}(N)} \leq 1 \quad [S127]$$

By substituting in the expression for $D_j(N)$ and using Stirling's approximation, we evaluate the limit on the left-hand side of Eq.S127 to express this inequality in terms of the parameters of the beta mixture distribution.

$$\lim_{N\to\infty} \frac{\omega_j B^{-1}(a_j, b_j) B(a_j, b_j + N)}{\omega_j B^{-1}(a_{j^*}, b_{j^*}) B(a_{j^*}, b_{j^*} + N)} \quad [S128]$$

$$= \frac{\omega_j \Gamma(a + b_j)\Gamma(b_{j^*})}{\omega_{j^*}\Gamma(b_j)\Gamma(a_{j^*} + b_{j^*})} \lim_{N\to\infty} \frac{\Gamma(b_j + N)\Gamma(a_{j^*} + b_{j^*} + N)}{\Gamma(a_j + b_j + N)\Gamma(b_{j^*} + N)} \quad [S129]$$

$$= C \cdot \lim_{N\to\infty} T_1 \cdot T_2 \cdot T_3 \quad [S130]$$

where

$$C = \frac{\omega_j \Gamma(a + b_j)\Gamma(b_{j^*})}{\omega_{j^*}\Gamma(b_j)\Gamma(a_{j^*} + b_{j^*})} \quad [S131]$$

$$T_1 = \sqrt{\frac{(b_j + N - 1)(a_{j^*} + b_{j^*} + N - 1)}{(a_j + b_j + N - 1)(b_{j^*} + N - 1)}} \quad [S132]$$

$$T_2 = \frac{e^{(a_j + b_j + N - 1) + (b_{j^*} + N - 1)}}{e^{(b_j + N - 1) + (a_{j^*} + b_{j^*} + N - 1)}} \quad [S133]$$

$$T_3 = \left(\frac{b_j + N - 1}{a_j + b_j + N - 1}\right)^{a_j + b_j + N - 1} \left(\frac{a_{j^*} + b_{j^*} + N - 1}{b_{j^*} + N - 1}\right)^{a_{j^*} + b_{j^*} + N} \frac{(b_{j^*} + N - 1)^{a_{j^*}}}{(b_j + N - 1)^{a_j}} \quad [S134]$$

We can evaluate the limits of $T_1$, $T_2$, $T_3$ using a similar approach to the steps taken in E. The limits of these three terms can be taken individually, where the steps to evaluate the limit of $T_1$ corresponds to the calculation of $\mathcal{L}_1$, and the same is true of $T_2$ and $\mathcal{L}_2$, and $T_3$ and $\mathcal{L}_3$ respectively. Applying these steps we get

$$\lim_{N\to\infty} T_1 = 1 \quad [S135]$$

$$\lim_{N\to\infty} T_2 = e^{a_j} e^{-a_{j^*}} \quad [S136]$$

$$\lim_{N\to\infty} T_3 = e^{-a_j} e^{a_{j^*}} \cdot t_3 \quad [S137]$$

$$t_3 = \lim_{N\to\infty} \frac{(b_{j^*} + N - 1)^{a_{j^*}}}{(b_j + N - 1)^{a_j}} \quad [S138]$$

The value of the limit in its entirety consists of the product of these terms, and therefore only depends on the values of $a_j$ and $a_{j^*}$ in the $t_3$ term. In particular, the limit is zero if $a_{j^*} < a_j$, undefined if $a_{j^*} > a_j$, and one if $a_{j^*} = a_j$. Therefore the inequality in Eq.S127 is always satisfied when $a_{j^*} < a_j$. Thus, for a significantly large $N$, we can define the index of the largest value of $D_j(N)$ such that $j^* = \{j \mid a_j = min(\hat{a}_j)\}$. Even when there is not a unique smallest value of $a_j$, any value in the set $j^*$ would provide almost equivalent upper bounds. To ensure the tightest possible upper and lower bounds it is necessary to find the unique value in $j^*$ that maximizes $D_j(N)$. This can be determined a priori, as this value only depends on the beta mixture parameters, but we find $D_j(N)$ will be maximized for the $j^*$ with the largest value of $b_{j^*}$. We then define this unique index $(\Omega)$ such that $D_\Omega \geq D_j$ for all values of $j$. This is reasonable as, decreasing the beta shape parameter $a$ and increasing the beta shape parameter $b$ both drive the mean of a beta distribution towards zero, which would have the greatest likelihood in a mixture of betas when the number of spillovers is large.

Using this observation, we define an upper and lower bound on the limit from Eq.S125 such that

$$\sum_{k=1}^{K} \lim_{N\to\infty} \frac{T_k(N)}{\sum_{j=1}^{K} D_\Omega(N)} \leq \sum_{k=1}^{K} \lim_{N\to\infty} \frac{T_k(N)}{\sum_{j=1}^{K} D_j(N)} \leq \sum_{k=1}^{K} \lim_{N\to\infty} \frac{T_k(N)}{D_\Omega(N)} \quad [S139]$$

For the upper bound term, we substitute the expressions for the numerator and denominator, so our upper bound limit can be expressed as

$$\sum_{k=1}^{K} \lim_{N\to\infty} \frac{\omega_k B^{-1}(a_k, b_k) B(a_k, b_k + (c+1)N)}{\sum_{j=1}^{K} \omega_j B^{-1}(a_j, b_j) B(a_j, b_j + N)} \leq \sum_{k=1}^{K} \lim_{N\to\infty} \frac{\omega_k B^{-1}(a_k, b_k) B(a_k, b_k + (c+1)N)}{\omega_\Omega B^{-1}(a_\Omega, b_\Omega) B(a_\Omega, b_\Omega + N)} \quad [S140]$$

DRAFT

Using the limit derivation from Eq.S138 and the fact that $a_\Omega \leq a_k$ for all values of $k$, we see that every limit term within the sum will be zero, except when $k \in j^*$, and applying these results, we can rewrite and compute the limit such that

$$\sum_{k=1}^{K} \lim_{N\to\infty} \frac{\omega_k B^{-1}(a_k, b_k) B(a_k, b_k + (c+1)N)}{\sum_{j=1}^{K} \omega_j B^{-1}(a_j, b_j) B(a_j, b_j + N)} \leq \frac{1}{(1+c)^{a_\Omega}} \sum_{k\in j^*} \frac{\omega_k \Gamma(a_k + b_k)\Gamma(b_\Omega)}{\omega_\Omega \Gamma(a_\Omega + b_\Omega)\Gamma(b_k)} \quad \text{[S141]}$$

The lower bound can be computed similarly, such that

$$\frac{1}{K \cdot (1+c)^{a_\Omega}} \sum_{k\in j^*} \frac{\omega_k \Gamma(a_k + b_k)\Gamma(b_\Omega)}{\omega_\Omega \Gamma(a_\Omega + b_\Omega)\Gamma(b_k)} \leq \sum_{k=1}^{K} \lim_{N\to\infty} \frac{\omega_k B^{-1}(a_k, b_k) B(a_k, B_k + (c+1)N)}{\sum_{j=1}^{K} \omega_j B^{-1}(a_j, b_j) B(a_j, b_j + N)} \quad \text{[S142]}$$

Thus, when there is a single smallest value of $a_j$, the lower and upper bounds are exactly equal. For this case, we have an exact analytical solution for the limit when the prior distribution follows a mixture of beta distributions, such that

$$P_\infty(H_F > 0|\pi(\phi)) = 1 - \frac{1}{(1+c)^{a_\Omega}} \quad \text{[S143]}$$

When there is not a unique smallest value of $a_j$, the lower and upper bounds will never be exactly equal, so we can only analytically compute bounds for the limit. Since we are interested in the probability of at least one host jump in the future, we only consider the upper bound on this probability as a conservative approach. To do this, we substitute our analytically computed lower bound, which gives

$$P_\infty(H_F > 0|\pi(\phi)) = 1 - \sum_{k=1}^{K} \lim_{N\to\infty} \frac{\omega_k B^{-1}(a_k, b_k) B(a_k, B_k + (c+1)N)}{\sum_{j=1}^{K} \omega_j B^{-1}(a_j, b_j) B(a_j, b_j + N)} \quad \text{[S144]}$$

$$P_\infty(H_F > 0|\pi(\phi)) \leq 1 - \frac{1}{K \cdot (1+c)^{a_\Omega}} \sum_{k\in j^*} \frac{\omega_k \Gamma(a_k + b_k)\Gamma(b_\Omega)}{\omega_\Omega \Gamma(a_\Omega + b_\Omega)\Gamma(b_k)} \quad \text{[S145]}$$

The gamma functions in this expression are strictly positive because parameters $\omega_k$, $a_k$, $b_k$, and $c$ are all greater than zero for all values of $k$, every term, we notice

$$\frac{1}{K \cdot (1+c)^{a_\Omega}} \sum_{k\in j^*} \frac{\omega_k \Gamma(a_k + b_k)\Gamma(b_\Omega)}{\omega_\Omega \Gamma(a_\Omega + b_\Omega)\Gamma(b_k)} > 0 \quad \text{[S146]}$$

Therefore, we have demonstrated that, when there is a linear correlation between the number of spillover events in the past and future, the probability of a host jump saturates to a value between zero and one as the rate of spillover approaches infinite values, and this is valid for any mixture of beta distributions. We find that this value depends heavily on the minimum value of the shape parameter $a_j$ in the mixture of beta distributions, although the other parameters are also relevant to the computed upper bound when the smallest $a_j$ is not unique.

**F. Supplementary Results.** To validate the analytical solution to our model, we use a stochastic model to simulate the probability of a future host jump ($H_F > 0$) for a given rate of spillover for the past ($\lambda$) and future ($c\lambda$), where there were exactly $H_P$ host jumps in the past. First, we simulate a random value of $\phi$ from our prior distribution $\pi(\phi)$, which we call $\phi_r$. We then use a Poisson distribution with mean $\lambda t_P$ to determine the number of past spillover events ($N$). Then, to determine the number of past host jumps ($H_P$), we use a binomial distribution with parameters ($n = N$, $p = \phi_r$), so each spillover had probability $\phi_r$ to result in a host jump. If this random value is exactly equal to the specified number of past host jumps ($H = H_P$), we store this value of $\phi_r$. Otherwise ($H \neq H_P$), the value of $\phi_r$ is discarded. We then simulate a new value of $\phi_r$ and repeat this process until we have stored 50,000 values on $\phi_r$. We then simulate the number of future host jumps ($H_F$) using a similar set of steps. We determine the number of future spillover events ($M$) using a Poisson distribution with mean $c\lambda t_F$. We then use a binomial distribution with parameters ($n = M$, $p = \phi_r$), to determine if a host jump will occur in $M$ future spillover events. We repeat this process with a new simulated value of $M$ for every stored value of $\phi_r$. The fraction of these binomial draws that did result in at least one host jump ($H_F > 0$) approximates the probability of a host jump given the model parameters $\lambda t_P$, $c\lambda t_F$, $H_P$, and $\pi(\phi)$. In figure S1 we see that the results of these simulations do indeed align with our analytical solution and produce the same qualitative trends suggested by our main results.

We also use our analytical solution for the count model derived in supplemental information C, with results shown in figure S2. We also validated these results using the simulation in figure S3, and we see that the simulation produces similar results to our analytical solution for the count-based model. Because time is implicit in the count model, it is not possible to replicate Fig2(J-L), but we see that this model still yields qualitatively similar results to the model presented in the main text, wherein the riskiest pathogens may be those that spill over at high, low, or intermediate rates.

**G. Evaluating reasonable prior hyper-parameters.** In the main text we chose parameters for our prior distributions to broadly characterize three biologically plausible scenarios. Here, we more carefully focus on defining the shape of our prior on $\phi$ by establishing reasonable values for the parameters, $a$ and $b$.

One possible approach to constructing a reasonable prior on $\phi$ would be to choose values for the beta shape parameters $a$ and $b$ such that the mean of the prior is equal to some estimate ($\bar{\phi}$). Host jumps are rare relative to the frequency of spillover events (18), implying that the mean of our prior should be small. A non-mixture beta distribution can take four general shapes, which arise under the following combinations of parameters $a$ and $b$:

1. $a,\ b \geq 1$
2. $a \geq 1,\ b < 1$
3. $a,\ b < 1$
4. $a < 1,\ b \geq 1$

In case 1, the prior is "hump-shaped", with a peak at some intermediate value between zero and one. Because we believe the mean of this distribution should be close to zero, the parameter $b$ must be much larger than $a$. Consequently, the variance will also be small, resulting in a prior that approaches a Dirac-delta function centered on $\bar{\phi}$ for large values of $b$. Such a prior would suggest that all pathogens are equally likely to host jump given a single spillover event. However, comparing human immunodeficiency virus and rabies lyssavirus illustrates a clear counterexample, suggesting that this scenario is unlikely to be realistic.

Case 2 produces a left-skewed distribution, wherein most pathogens have host jump probability close to one. Rationally this can not be the case, as most spillover events do not result in host jumps. Therefore, between cases 1 and 2, we have established that the parameter $a$ must be less than one.

Case 3 generates a prior distribution that is ”U-shaped”, and the parameter $a$ must be much smaller that $b$ in order to have a sufficiently small mean. However, this prior suggests that there are some pathogens that will almost certainly host jump given a spillover event. Indeed, some pathogens are much more likely to host jump than others, but stochastic effects and other demographic complexities make such high host jump probabilities unlikely, even for pathogens with a large $R_0$.

In case 4, the prior is right-skewed, such that most pathogens are unlikely to complete a host jump. Because host jumps are thought to be rare, stochastic events, this would seem like the most reasonable shape for a prior using a single beta distribution. In this case, the suspected small mean value of $\phi$ can be achieved by decreasing the value of $a$ or increasing the value of $b$. However, there are multiple combinations of $a$ and $b$ that produce equal means, so we further evaluate how changing these parameter values while retaining a constant mean would affect our conclusions.

Lastly, we note that the observations for these four cases do not necessarily apply to their usage in a mixture of beta mixture distributions, as different combinations may be able to better capture variation between different families of pathogens that a single beta distribution alone can not. However, as we have shown in Supplementary text D2, the limiting behavior of our model when using a beta mixture prior depends only on the beta component that has the smallest value for the parameter $a$. Thus even when a beta mixture contains a beta from cases 1-3, its behavior as spillover rate goes to infinity will be determined by the behavior of case 4.

***2. Results under changing prior hyper-parameters.*** Since a prior distribution with a reasonable mean can be achieved by changing both $a$ and $b$, we would like to characterize the effects of changing the values of these parameters. As we demonstrated in Supplementary text E, the probability of a host jump converges to a value between zero and one, which only depends on the parameter $a$, the slope $c$ of the linear relationship between the number past and future spillover events, and the past and future time horizons ($t_P$ and $t_F$). Therefore, as the rate of spillover gets large, decreasing the value of $a$ will result in a decrease in the probability of a host jump, but decreases in the value of $b$ will have no impact on that probability. However, as we show below, increasing the value of $b$ decreases the rate at which convergence is achieved. We show the difference between changing either $a$ or $b$ in figure S4.

We also quantify the rate of convergence using a standardized distance metric under different values of $a$ and $b$, shown in figure S5. We define this distance metric metric as:

$$Std\_Dist = \frac{|\mathcal{F}(N) - \mathcal{L}|}{\mathcal{L}} \qquad [S147]$$

where $\mathcal{F}(N)$ is the analytical solution to our model along the $c:1$ line, and $\mathcal{L}$ is the value of the limit derived in D. In general, we see that this metric is relatively insensitive to changes to the parameters $a$ and $c$ for any given scenario; however, we do see that this observation does not hold for large values of $b$. In scenarios 1 and 3 in figure S5**B**, we see that increasing values of $b$ leads to larger values for our distance metric, implying that convergence is slower for large values of $b$. From this figure, we also recognize that increasing $a$ or $c$ to comparably large values would not give the same results, since our distance metric decreases as $a$ or $c$ are increased, and the rate of convergence metric is almost completely insensitive to changes in these parameters.

We also demonstrate the effects of changing the values of $a$ and $b$ in while retaining a fixed mean for the prior distribution. We focus specifically on the right-skewed prior (similar to scenario 1 in the main text) and the right-skewed component of the beta mixture distribution (similar to scenario 3 in the main text). We define a conservative hypothetical mean for our prior such that $\bar{\phi} = 0.01$, and restrict the prior parameters such that $a < 1$ and $b \geq 1$. We use three sets of parameters to broadly characterize possible results within this parameter space and define our three right skewed priors as:

$$\pi(\phi) \sim \text{Beta}(a = 0.0102,\ b = 1.0098) \qquad [S148]$$

$$\pi(\phi) \sim \text{Beta}(a = 0.9999,\ b = 98.9901) \qquad [S149]$$

$$\pi(\phi) \sim \text{Beta}(a = 0.3,\ b = 29.7) \qquad [S150]$$

In addition to the single beta distributions, we also use these parameters in a mixture distribution, similar to Scenario 3 in the main text, where our prior has the form $\pi(\phi) \sim 0.8 \cdot \text{Beta}(a,\ b) + 0.2 \cdot \text{Beta}(100,\ 400)$, where the first component in the mixture is replaced with each of the three aforementioned right-skewed beta distributions.

We show our model predictions under these different prior parameterizations in figure S6. We see that pathogens that frequently spill over pose the greatest host jump risk for each of these priors. As expected from the results in figure S4, the magnitude of this risk depends on the value of $a$, and the rate at which this level of risk is achieved depends on $b$. While the conclusion as to which types of pathogens are most likely to host jump is the same for each of these priors, the magnitude of this risk is fundamentally different. While all three priors have the same mean, this dramatic difference in the magnitude of risk posed by zoonotic pathogens highlights the importance of our ability to estimate meaningful values for our prior parameters.

Similarly, we also show our model predictions for several beta mixture distributions with different parameter values for the right-skewed component of the mixture in figure S7. Here, we see that these different parameterizations may lead to different conclusions as to which pathogens are most likely to host jump. When the value of the parameter $a$ is small, we see that host jump risk is highest for intermediate levels of spillover due to the subset of ”higher-risk pathogens” represented by the second component of the mixture. However, when the value of $a$ is sufficiently large in the right-skewed component, the subset of ”high risk pathogens” characterized by the second mixture component are ”out-competed” by pathogens that frequently spill over in terms of host jump risk. Therefore, pathogens that frequently spill over are most likely to host hump in these cases.

Based on these results for different prior parameterizations, we see that our ability to evaluate the relationship between spillover and host jump risk is dependent on prior distribution, or rather, our ability to quantify our uncertainty on the value of $\phi$. We see that our choices for the values of $a$ and $b$ play a significant role in quantifying the risk posed by zoonotic pathogens. Additionally, we see that the components of our mixture distribution determine whether potential high-risk pathogens that rarely spill over are more likely to host jump than pathogens that frequently spill over. The shape of this distribution (and the parameters that define it) will depend heavily on our understanding of characteristics of pathogens, the hosts, and the environment that either facilitate or hinder the process of a successful host jump. Past and future research on these subjects will be critical to characterizing a prior on $\phi$ that allows us to accurately evaluate the relationship between spillover and host jump risk and identify contexts where host jumps are most likely.

DRAFT

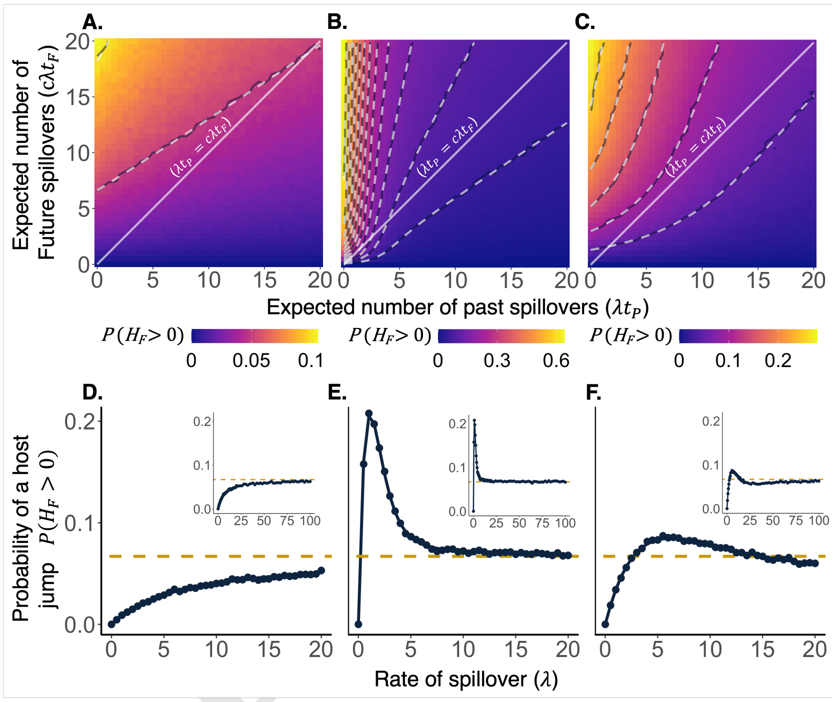


**Fig. S1.** When the number of spillover events in the past and future follow independent Poisson distributions, our model results are qualitatively similar to the results found in the main text. Panels A-C show the probability of a future host jump for each of our three scenarios from the main text when the number of past and future spillover events follow Poisson distributions with fixed rates $\lambda$ and $c\lambda$ respectively. We generate these probabilities using 50,000 replicates in our simulation, and the contour lines are shown for every 5% increment of host jump risk for the simulation (solid line) and our derived numerical solution (dashed line). In panels D-F, we show host jump risk when past and future rate parameters are linearly correlated for both the simulation (points) and our derived numerical solution (solid line). These trends are qualitatively similar to the results shown in the main text. It appears that very high rates of spillover may approach the same value derived in E (dashed gold line).

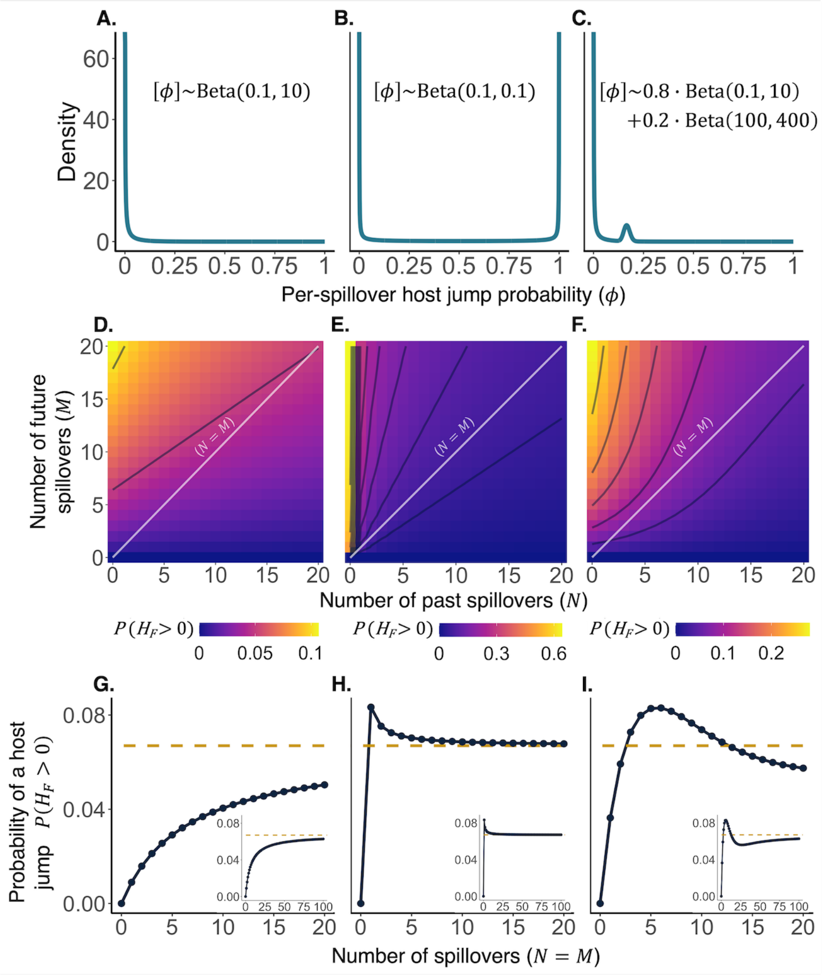


**Fig. S2.** The columns of panels from left to right denote Scenarios 1-3 respectively. Panels A-C show the prior distributions on $\phi$. Panels D-F show the model-calculated probability of a successful host jump as a function of the number of past and future spillover events. Host jump risk increases with future spillovers (M) and decreases with past spillover (N). The contour lines in each panel demarcate changes of 5% in total host jump risk. Note that the color range differs for each panel. The solid white lines represent the 1:1 relationship between the number of past and future spillovers, and panels G-I show the probabilities of a future host jump along these lines. Comparing panels G-I reveals that depending on $[\phi]$, host jump risk may increase or decrease as a function of spillover rate. Nevertheless, in all scenarios, host jump risk converges to the value $\left(1-\frac{1}{(c+1)^{a}}\right)$, as spillover events increase (inset figures).

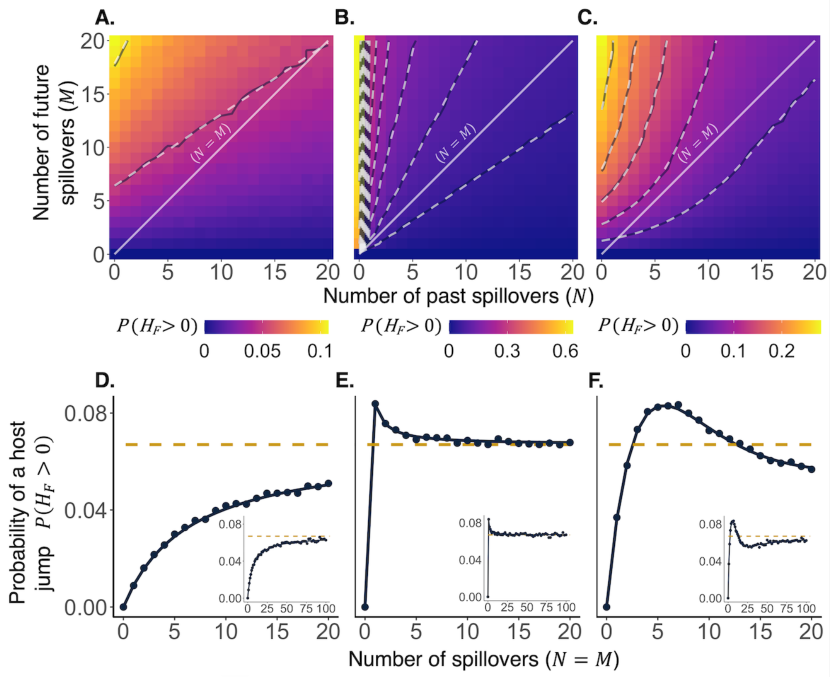


**Fig. S3.** Panels A-C indicate the fraction of 50,000 simulations where a host jump occured for the given number of past (N) and future (M) spillover events. Contour lines representing a 5% change in host jump risk are shown for the simulation values (solid lines), which closely follow the analytic solution (dashed lines). Panels D-F show the analytical solution from the main text (line) with simulation results (points) when past and future are linearly correlated, and confirm the results in the main text.

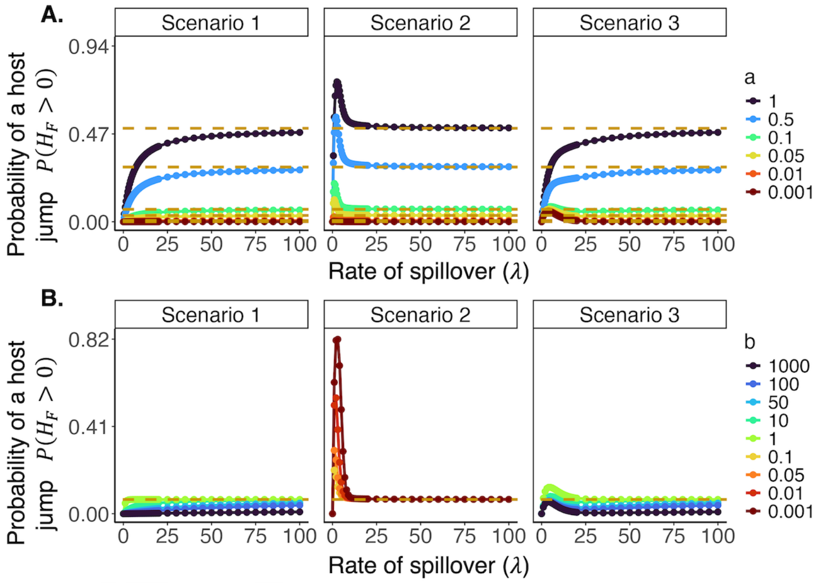


**Fig. S4.** Here we show the effects of changing the values of different prior hyper-parameters in the three scenarios discussed in the main text. Results for different values of the parameter $a$ are shown in panel **A** and results for different values of the parameter $b$ are shown in panel **B**. For the beta mixture distribution, only the shape parameters in the first term of the mixture (i.e., $0.8 \cdot \text{Beta}(a = 0.1,\ b = 10)$) are changed. In panel **A**, we see that changing the value of $a$ affects the limit value in each of these scenarios, but all curves converge at approximately similar rates. In contrast, we see that changing the value of $b$ in panel **B** does not change the limit value, but the curves take longer to saturate for large values of $b$. Note that not all values of $b$ are used in each scenario in panel **B**. Scenarios 1 and 3 do not use $b < 1$ to avoid redundancy with scenario 2. Similarly, scenario 2 does not use $b \geq 1$, as this would yield a left-skewed prior, which is not biologically realistic.

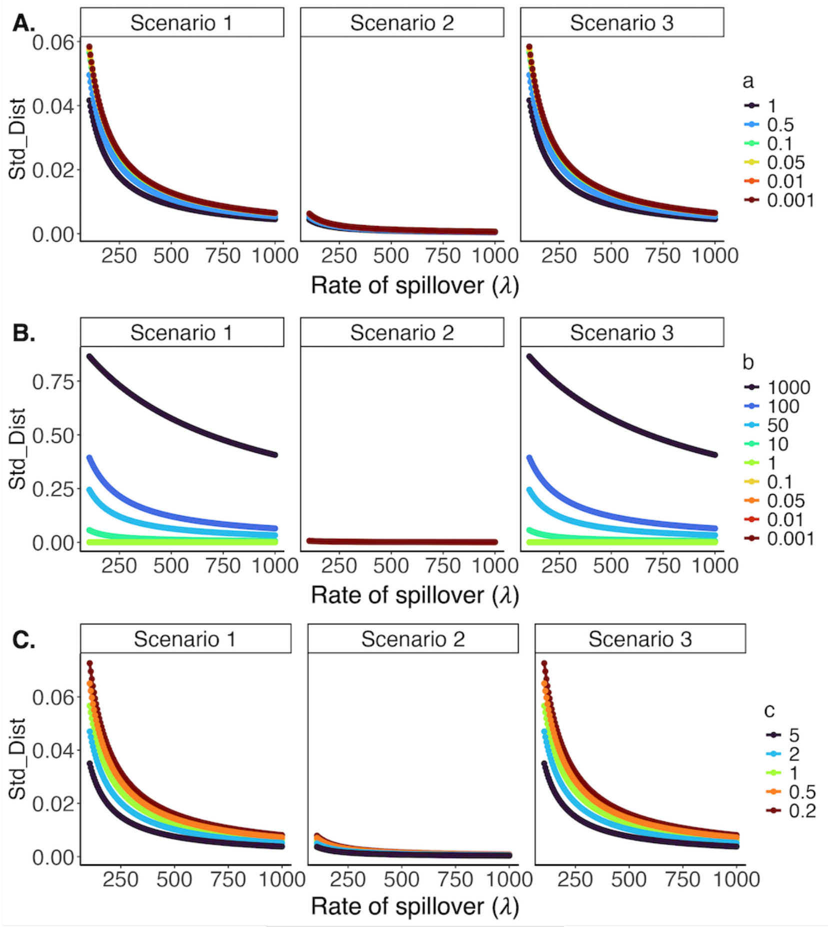


**Fig. S5.** Here we show how changing the values of our model parameters in the three scenarios from the main text affect the relative distance to convergence. Results are shown for different values of $a$ (panel **A**), $b$ (panel **B**), and $c$ (panel **C**). For the beta mixture distribution in scenario 3, only the shape parameters in the first term of the mixture (i.e., $0.8 \cdot \text{Beta}(a = 0.1,\ b = 10)$) are changed. We see that the distance to convergence is relatively insensitive to changes in the value of parameters $a$ and $c$. However, this distance is highly sensitive to increasing values of $b$, as our standardized distance metric is greatest when $b$ is large. Importantly, we note that this same trend is unlikely to hold when increasing $a$ or $c$, as we see here that increasing these values leads to faster convergence.

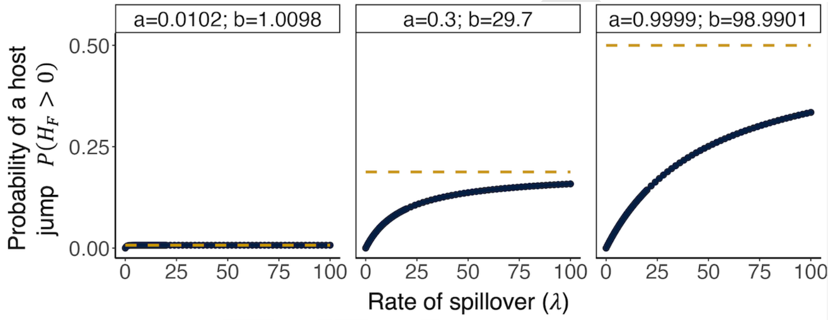


**Fig. S6.** Here we show how changing the values of the prior hyper-parameters for a right-skewed prior (like scenario 1) would affect our results. In all cases, we see that these curves all eventually converge to a value between zero and on depending on the value of $a$, and that the speed at which they converge to this value depends on $b$. Parameters are chosen such that all three priors used here have equal means of $\bar{\phi} = 0.01$. When $a$ is small, $b$ must be close to one, so the probability of a future host jump converges to a small value quickly. As the values of $a$ and $b$ increase, the probability of a future host jump increases while the rate of convergence decreases. Similar to our results for a right-skewed prior in the main text (Scenario 1), we see that host jump risk is strictly increasing with respect to the number of spillover events.

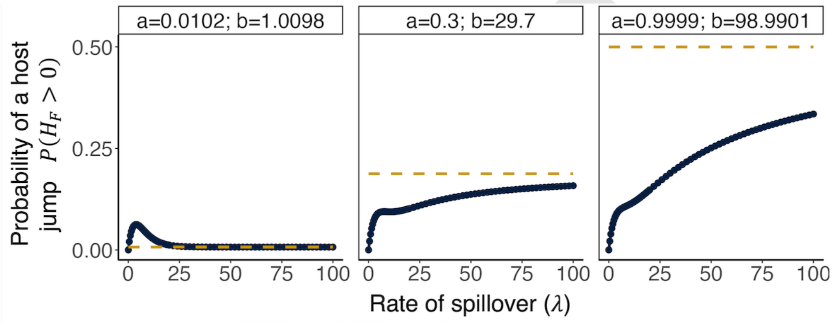


**Fig. S7.** Here we show how changing the values of the prior hyper-parameters for the right-skewed component of our beta mixture prior (like scenario 3) would affect our results. We still see that the parameters of the right skewed component control the value and rate of saturation. However, unlike our results for the mixture distribution in the main text (Scenario 3), it is not always the case that that intermediate levels of spillover pose the greatest host jump risk. We see that this maximum risk at intermediate levels of spillover also depends on the values of $a$ and $b$ in the right-skewed component of the mixture distribution. Instead, when the parameter $a$ is sufficiently large, pathogens that spill over most pose the greatest host jump risk.
